# The β integrin modulates serotonin sensitivity via NPxY motifs to regulate egg laying and mechanosensation behaviors in *C. elegans*

**DOI:** 10.1101/2024.12.23.630109

**Authors:** Josh Haram Bumm, Daniel Dogeon Lee, Zhongqiang Qiu, Angela Haeun Choi, Meghana Kolluri, Michael J. Olsen, Lianzijun Wang, Myeongwoo Lee

**Affiliations:** Department of Biology, Baylor University, Waco, Texas 76798, USA

**Keywords:** phosphomimetic, serotonin transporter, fluoxetine, reserpine, talin, MEC-4. Running title: Role of integrin NPxY motifs in egg laying and mechanosensation

## Abstract

**Background:** Integrin is an αβ heterodimeric receptor to the extracellular matrix; its binding to the matrix recruits focal adhesions to two NPxY motifs, the tyrosine phosphorylation sites in the cytoplasmic domain. Studies found that replacing tyrosines (Y) with phenylalanines (F) in the motif of β1 integrin displayed little developmental or behavioral defects. However, the tyrosine-to-alanine (A) caused embryonic lethality.

**Results:** Here we report novel functions of the NPxY motifs in *C. elegans pat-3* β integrin. The membrane-proximal non-phosphorylation *pat-3(Y792F)* mutation caused hypersensitive egg laying in serotonin, which is more prominent than the membrane-distal NPxY^804^. The double non-phosphorylatable *pat-3(YYFF)* mutant exhibited serotonin hypersensitivity and defective egg retention. The phosphomimetic NPxY, *pat-3(Y804E),* mutant displayed a reduced egg laying in response to serotonin and fluoxetine, suggesting that the NPxY phosphorylation is associated with vulval contraction and serotonin sensitivity. Additionally, *pat-3(Y792A)*, *pat-3(Y792F)*, *pat-3(Y804E), and pat-3(YYFF)* mutants exhibited mechanosensation defects, demonstrating that NPxY phosphorylation regulates sensory neuron activity. Further revealed that exogenous serotonin reduced mechanosensation, while blocking serotonin secretion rescued the mechanosensation of *pat-3* NPxY mutants, suggesting that integrin NPxY modulates serotonin levels in *C. elegans*.

**Conclusion:** Our results underscore the functional importance of *pat-3* NPxY motifs in muscle and neurons, potentially linking integrin NPxY motifs to neurotransmitter response and mechanosensory functions.

## INTRODUCTION

Integrin is an αβ heterodimeric cell surface receptor to the extracellular matrix (ECM), controlling cell behaviors through bidirectional signaling. Inside-out signaling activates the integrin extracellular domain to facilitate ECM binding, while outside-in recruits signaling molecules and links cytoskeletons to the cell-ECM contacts, known as focal adhesions (FAs) ^1^. In this context, the cytoplasmic domain (CTD) of β integrin interacts with linker proteins such as talins and kindlins to transduce mechanical information from the ECM to the cell ^2^. Two conserved NPxY (Asn-Pro-x-Tyr) motifs, potential tyrosine phosphorylation sites, exist in the CTD of all metazoan β integrins ^3^ (Figure 1). The membrane-proximal NPxY binds to the FERM domain of talin, while the membrane distal motif interacts with kindlin ^4^. Sakai *et al.* reported overexpression of *src* kinase in a mouse fibroblast phosphorylated the NPxY motifs of β1A integrin ^5^. In mice, knock-in β1 integrin YYFF (Tyrosine-to-Phenylalanine) failed to exhibit discernible phenotypes ^6^, although integrin NPxY mutants showed resistance to tumorigenesis ^7^ and defects in blood clotting ^8^. Conversely, disruptive (tyrosine removal) β1 YYAA, Y783A, or Y795A (Tyrosine-to-Alanine) mutations caused severe phenotypes such as embryonic lethality ^6^. These findings suggested that the Y-to-A transition generally brings severe phenotypes to the animal, while the Y-to-F transition produces no overt developmental phenotypes.

**Figure 1.**
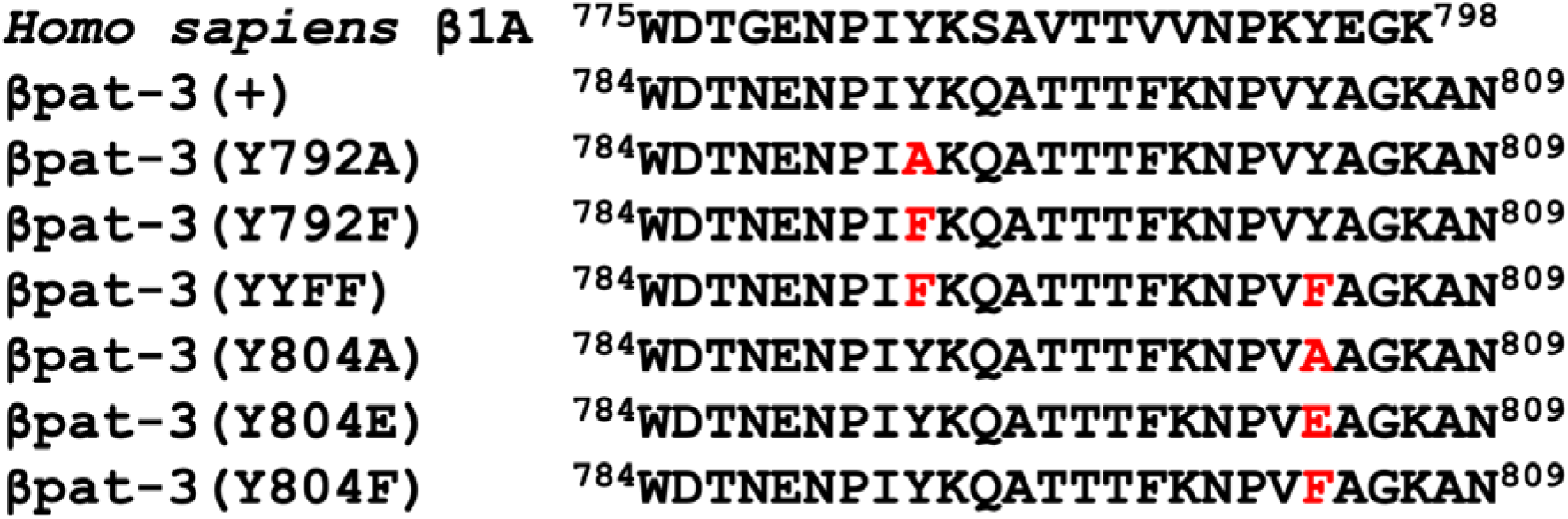
Amino acid sequences of wild-type (+) and mutant cytoplasmic domain (CTD) of *C. elegans* βpat-3 integrin. Amino acid residues in *C. elegans* βPAT-3 cytoplasmic tail sequence and their functions. The amino acid sequence of the *Homo sapiens* β1A integrin CTD is aligned with *C. elegans pat-3 (+),* wild-type CTD, and mutant CTD such as *pat-3(kq7921, Y792A), pat-3(kq7923, Y792F), pat-3(kq24, YYFF), pat-3(kq8041, Y804A), pat-3(kq8042, Y804E), and pat-3(kq8043, Y804F).* Reds indicate mutations.

In the nematode *C. elegans*, PAT-3/β integrin is ubiquitously localized in tissues, including body wall muscles, gonads, neurons, vulva, and the excretory canal ^9^. PAT-3 integrin also contains two NPxY motifs ^10^. Previous analyses by Itoh *et al.* suggested that SRC family tyrosine kinases (SFK) could phosphorylate the *pat-3* CTD. For instance, *C. elegans src-1* mutant exhibited gonad phenotypes similar to those caused by the integrin HA (hemagglutin)-βtail dominant negative allele ^10,11^. Our recent study demonstrated that mutations in the membrane distal Y to A, E (glutamic acid), or F mutations resulted in gonad migration and mild motility impairments ^12^. These findings suggest that the membrane-distal NPxY is dispensable for development and motility. However, given the widespread distribution of integrins in muscle, neurons, and gonads, we hypothesized that the NPxY motifs in PAT-3 CTD are crucial in undescribed tissue-specific functions.

In *C. elegans* hermaphrodites, vulva development begins at the L1 larval stage and culminates at the late larval stages. In the early adult stages, fertilized eggs accumulate in the uterus. Within eight hours of post-fertilization, the fertilized eggs are expelled from the uterus through vulva contractions, known as ‘egg laying,’ which is regulated by a pair of serotonergic hermaphrodite-specific motor neurons (HSNs) connected to vulva muscles. During egg laying, the HSN releases a neurotransmitter, serotonin, onto the vulva muscle, inducing its contractions. When the serotonin production or its receptor is defective, the vulva muscles fail to contract, accumulating unlaid eggs in the uterus and causing the Egl (<u>eg</u>g-laying defective) phenotype. Treating worms with exogenous serotonin or selective serotonin reuptake inhibitors (SSRIs) stimulates vulva muscle contraction and increases egg laying ^13^. Mutations in the genes associated with muscle attachment ^14^, such as βPAT-3 RNAi ^13,15^, perlecan/UNC-52 ^16,17^, and syndecan/SDN-1 ^18^, have been reported to cause defective egg laying, suggesting a link between the Egl phenotype and impaired cell-ECM interactions.

In *C. elegans*, gentle touch on the body induces backward or forward movement. The touch stimulus is detected by six touch receptor neurons (TRNs): ALMR, ALML, AVM, PLMR, PLML, and PVM ^19^. These TRNs express the MEC-4 DEG/ENaC channel complex on their cell surfaces, transmitting touch stimuli to the neurons. Extensive studies have identified several *mec* (mechanosensation abnormal) genes essential for processing touch-induced behaviors ^20^. Integrins and focal adhesion proteins transmit mechanical forces and trigger cellular signaling responses ^21^. Focal adhesion proteins are expressed in mechanosensory cells, including TRNs ^22^. Studies on focal adhesion genes—such as *unc-97/PINCH* ^23^ mutant and removal of *pat-2*/α integrin ^24^, *pat-3*/β integrin ^10^, *pat-6*/actopaxin ^25^, and *unc-112*/kindlin ^26^—in ALM and PLM touch neurons of genetic mosaic animals demonstrated the involvement of these genes in mechanosensation ^22^.

In this study, our results revealed that the phosphomimetic *pat-3 (kq8042, Y804E)* mutation exhibited defective egg-laying in response to serotonin or fluoxetine treatment. Conversely, the membrane-proximal non-phosphorylation *pat-3 (kq7923, Y792F)* mutant was hypersensitive to serotonin, while the membrane-distal NPxY mutations showed varied serotonin hypersensitivity. In contrast, the double non-phosphorylation *pat-3(kq24, YYFF)* mutant displayed a wild-type response to serotonin hypersensitivity and increased egg retention in the uterus. Further investigation into the role of *pat-3* NPxY motifs revealed that *pat-3(kq7921, Y792A)*, *pat-3(kq7923, Y792F)*, *pat-3(kq24, YYFF)*, and *pat-3(kq8042, Y804E)* mutants exhibited touch insensitivity in the anterior or posterior regions of the body. Subsequent analysis revealed that exogenous serotonin reduced mechanosensation, while inhibiting neurotransmitter secretion by reserpine, a vesicular monoamine transporter (VMAT) inhibitor ^27^, rescued the mechanosensation of *pat-3* NPxY mutants. These findings suggest that phosphorylation of the NPxY motif is critical for modulating serotonin response in both egg-laying machinery and mechanosensation.

## RESULTS

### Mutations in the βpat-3 NPxY motifs

To characterize the phosphorylation of the NPxY motif in the β*pat-3* integrin CTD, we introduced mutations into the membrane-distal NPxY in the β*pat-3* CTD (Figure 1). Previous analyses revealed that CRISPR edits in the membrane-distal NPxY of β*pat-3* led to motility and gonad migration defects ^12^. Additionally, we generated *pat-3(kq24, YYFF)*, a non-phosphorylatable mutation. Table 1 summarizes the behavioral phenotypes of *pat-3* NPxY mutants described in this manuscript.

**Table 1.**
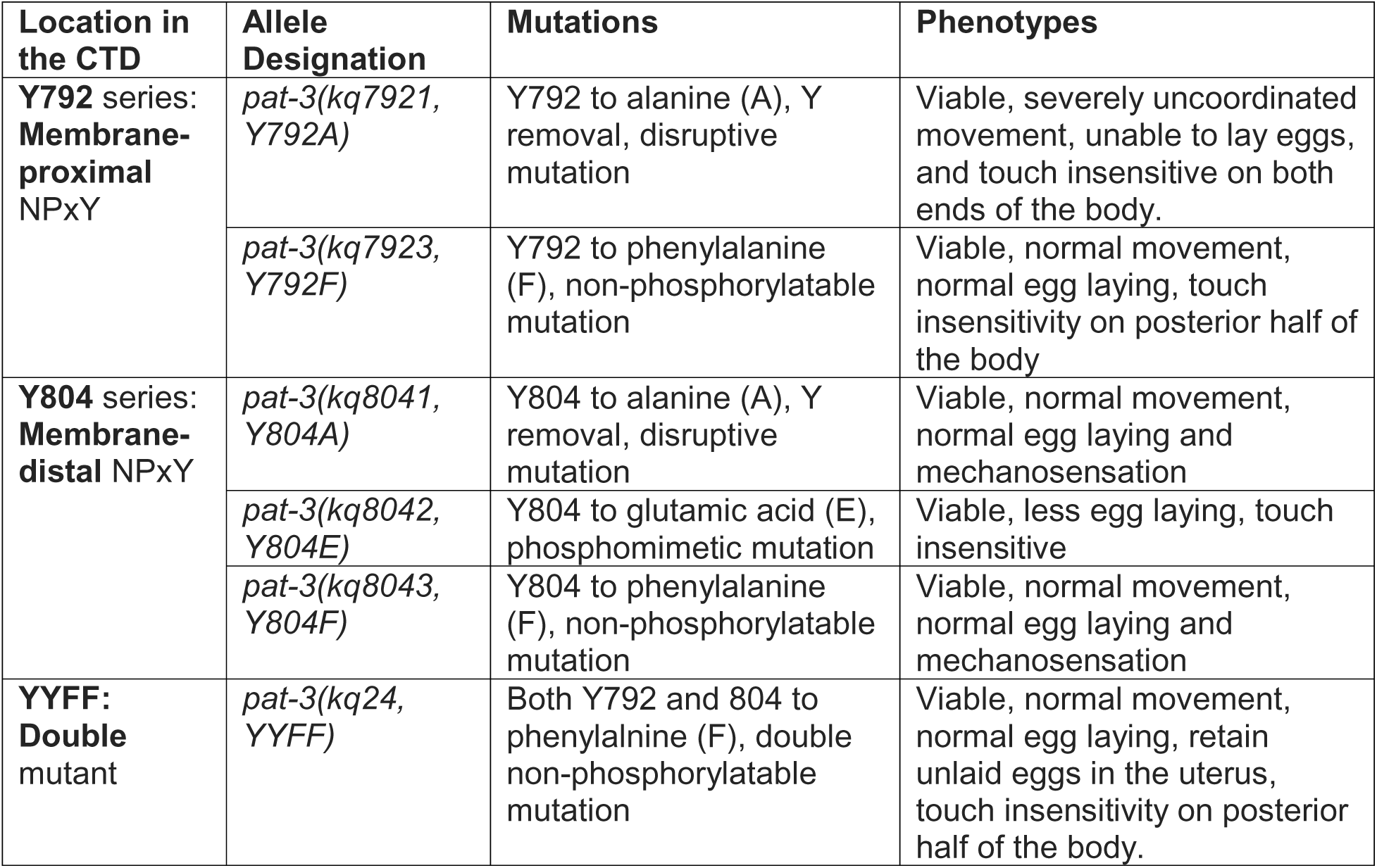
Summary of *pat-3* NPxY mutant phenotypes.

| Location in the CTD | Allele Designation | Mutations | Phenotypes |
| --- | --- | --- | --- |
| **Y792 series:**<br>**Membrane-proximal**<br>NPxY | *pat-3(kq7921, Y792A)* | Y792 to alanine (A), Y removal, disruptive mutation | Viable, severely uncoordinated movement, unable to lay eggs, and touch insensitive on both ends of the body. |
|  | *pat-3(kq7923, Y792F)* | Y792 to phenylalanine (F), non-phosphorylatable mutation | Viable, normal movement, normal egg laying, touch insensitivity on posterior half of the body |
| **Y804 series:**<br>**Membrane-distal** NPxY | *pat-3(kq8041, Y804A)* | Y804 to alanine (A), Y removal, disruptive mutation | Viable, normal movement, normal egg laying and mechanosensation |
|  | *pat-3(kq8042, Y804E)* | Y804 to glutamic acid (E), phosphomimetic mutation | Viable, less egg laying, touch insensitive |
|  | *pat-3(kq8043, Y804F)* | Y804 to phenylalanine (F), non-phosphorylatable mutation | Viable, normal movement, normal egg laying and mechanosensation |
| **YYFF:**<br>**Double**<br>mutant | *pat-3(kq24, YYFF)* | Both Y792 and 804 to phenylalanine (F), double non-phosphorylatable mutation | Viable, normal movement, normal egg laying, retain unlaid eggs in the uterus, touch insensitivity on posterior half of the body. |

Most of the *pat-3* NPxY mutants exhibited regular muscle filament organization. Some mutants were crossed with AH3437 *GFP::tln-1*, a GFP inserted in the N-terminus of TLN-1 using the CRISPR-Cas9 system ^28^, to visualize the structure of dense bodies and M lines (Figure 2, panels A to F). TLN-1 is expressed in body wall muscle and gonad epithelial cells, displaying typical dense bodies and M-line localization (arrows in Figure 2). Except for *pat-3(kq7921, Y792A)* in Figure 2E, all other mutants displayed intact dense bodies along the body wall, which is similar to that of AH3437 GFP::TLN-1 (Figure 2A), suggesting that the mutations were dispensable for organizing cell attachment structures to the ECM. In addition to GFP::TLN-1, the NPxY mutants were stained with rhodamine-conjugated phalloidin (Figure S1). Other than the deformity of muscle filaments caused by the staining procedures, we failed to detect an abnormal pattern of actin filaments.

**Figure 2.**
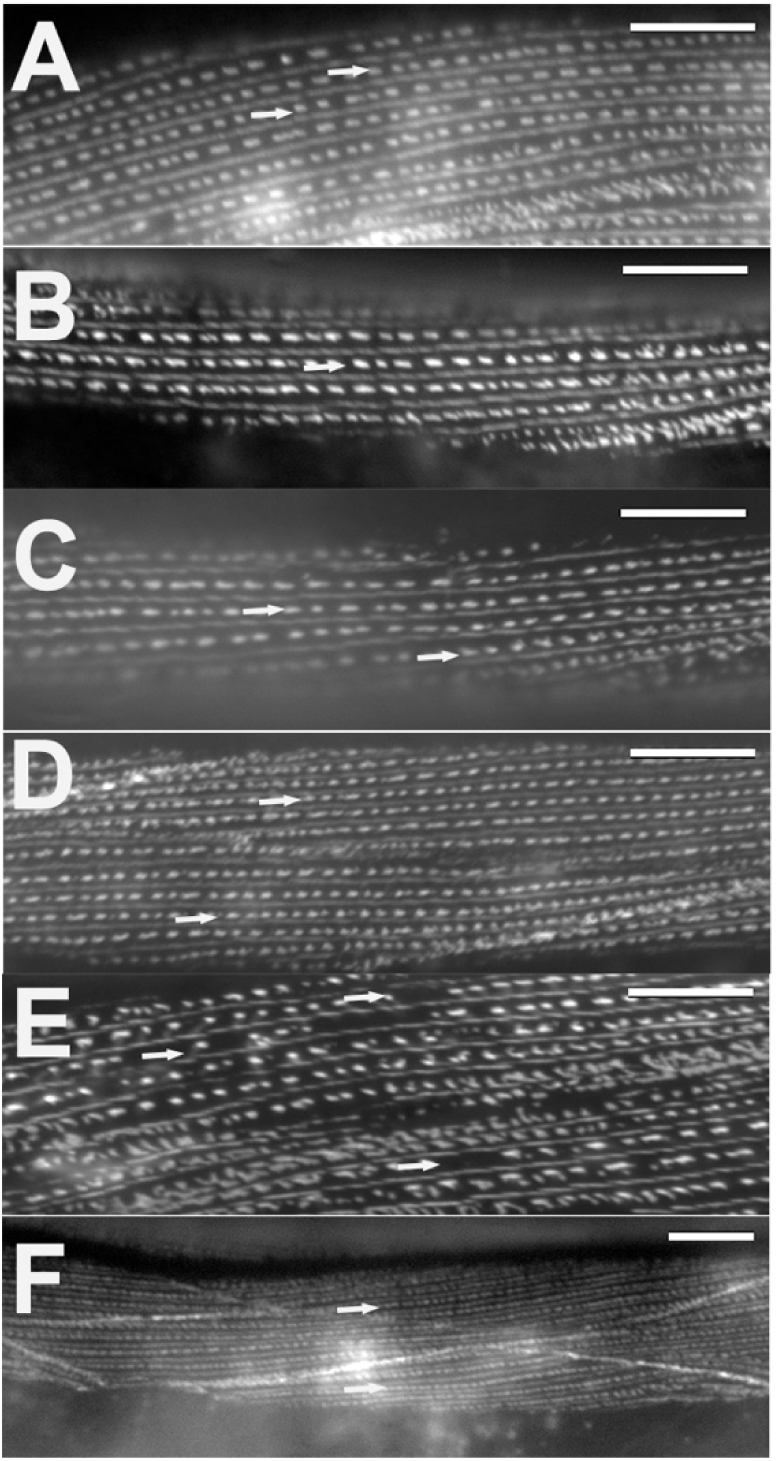
*GFP::tln-1/talin* localization in *pat-3* NPxY mutants The five double mutants were generated by crossing GFP::*tln-1* to *pat-3* NPxY mutants. In every picture, horizontal dots represent dense bodies (arrows), and thin lines between the dense bodies represent M lines ^60^. All images are taken under a 60X objective lens, except panel F (40X). Homozygous mutants are shown here. A. AH3437 *GFP::tln-1/talin* body wall muscle of adult worm (control), B. *GFP::tln-1/talin; pat-3(kq8042, Y804E),* C. *GFP::tln-1/talin; pat-3(kq8041, Y804A),* D. *GFP::tln-1/talin;pat-3(kq7923, Y792F),* E. *GFP::tln-1/talin;pat-3(kq7921, Y792A),* F. *GFP::tln-1/talin ;pat-3(kq24, YYFF)*. Bars = 10 μm.

### The membrane-proximal and distal NPxY motifs affect egg-laying behavior

Although the non-phosphorylatable NPxY lacks distinct phenotypes in mammalian β integrins ^6^, the importance and conservation of the β integrin NPxY led us to investigate the undescribed role of NPxY in integrin functions. Therefore, we sought to discover functional phenotypes associated with the NPxY mutations. When the hermaphrodite is submerged in an aqueous solution, it pauses egg laying because worms recognize the environment as a lack of food ^29^. Adding exogenous serotonin would stimulate vulva contraction, which results in egg laying ^13^. Before egg-laying assays, the fecundity of viable NPxY mutants, N2, and other control mutants was assessed to determine the optimal stages for measuring egg laying (Figure S2). All worm strains showed the peak number of eggs on Day 2 (see Experimental Procedures). Therefore, the egg-laying behavior was assessed with Day 2 hermaphrodites.

To monitor the egg laying of *pat-3* NPxY mutants, we incubated Day 2 adult hermaphrodites in 3.0 mg/ml (17 mM) serotonin dissolved in M9 buffer. After one hour of incubation, *pat-3(kq8042, Y804E)*, the phosphomimetic mutant, laid significantly fewer eggs in the serotonin solution than N2 (*p < 0.0001*). The *pat-3(kq8043, Y804F),* the distal non-phosphorylatable, and *pat-3(kq8041, Y804A)*, the distal tyrosine removal, mutants showed comparable egg laying to N2 (*p < 0.8*) (Figure 3, Supplementary Table 3).

**Figure 3.**
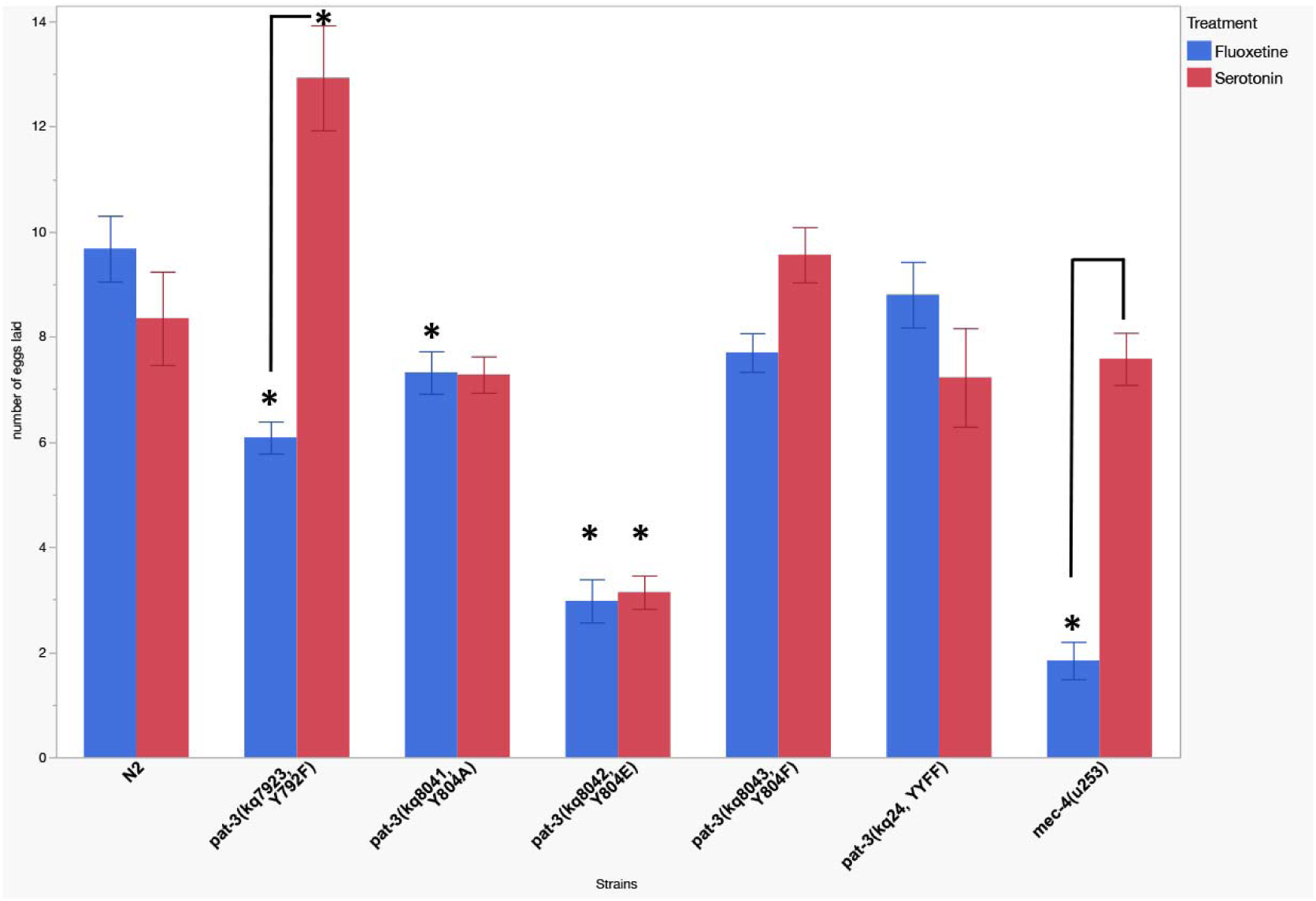
Egg-laying assays of integrin mutants. Results significantly different from N2 were marked with * (*p* < 0.05). Brackets indicate significant differences between the treatments. Multiple adult worms (n=40 or 50) were used for each experiment. *All Pairwise Comparisons - Tukey HSD.

Active egg laying is stimulated by a pair of serotonergic HSN motor neurons, which secrete serotonin (5-hydroxytryptamine, 5-HT) to induce vulva muscle contraction. Secreted serotonin is then removed by the serotonin transporter (SERT), which is expressed on the vulval tissue, as it enters inactive egg laying ^30^. Previous studies demonstrated that selective serotonin reuptake inhibitors (SSRIs) such as fluoxetine block the function of SERT. It is suggested that fluoxetine binds to MOD-5/CeSERT and stimulates *C. elegans* egg laying by blocking serotonin removal in the vulval tissue ^31^.

To test whether the Egl of *pat-3* NPxY mutants was due to abnormal 5-HT secretion, wild-type N2 and the mutants were treated with 0.5 mg/ml (1.6 mM) of fluoxetine ^32^. In N2, fluoxetine treatment would yield comparable egg-laying results to serotonin treatment (Figure 3). However, the egg laying of *pat-3(Y804E)* in fluoxetine was defective (*p < 0.0001*), suggesting that the reduced egg laying of the mutant was due to defective vulva contractions (Figure 3, Supplementary Table 3). The membrane-proximal *pat-3(kq7923, Y792F)* was also generated to compare with the membrane-distal NPxY mutants. The *pat-3(Y792F)* worms showed a significantly increased egg-laying response to serotonin compared to N2 (mean = 12.8 eggs/hour, *p < 0.0001,* Figure 3).

### The membrane-proximal NPxY mutant showed hypersensitivity to serotonin and fluoxetine

Our results suggested that the proximal non-phosphorylation, *pat-3(Y792F),* increased the response to exogenous serotonin, resulting in hyperactive egg laying (Figure 3). To further analyze the increased egg laying in *pat-3(Y792F)*, we diluted serotonin to a concentration that resulted in no N2 egg laying. In Figure 4, N2 animals incubated in 0.1 mg/ml (565 μM) serotonin could not stimulate egg laying. In contrast, *pat-3(Y792F)* showed a significant egg-laying increase at the diluted serotonin concentration (565 μM). At the same time, the number of eggs laid was comparable to N2 in 3 mg/ml serotonin (Figure 3), suggesting that the proximal non-phosphorylation would cause the hypersensitive egg laying in serotonin and that the proximal NPxY regulates the serotonin sensitivity. *pat-3(Y792F)* also displayed egg laying in 0.1 mg/ml fluoxetine, a concentration that failed to stimulate egg laying in N2 (Figure 4), suggesting this mutant is hypersensitive to fluoxetine treatment, although the response was significantly lower than that of serotonin at the diluted concentration.

**Figure 4.**
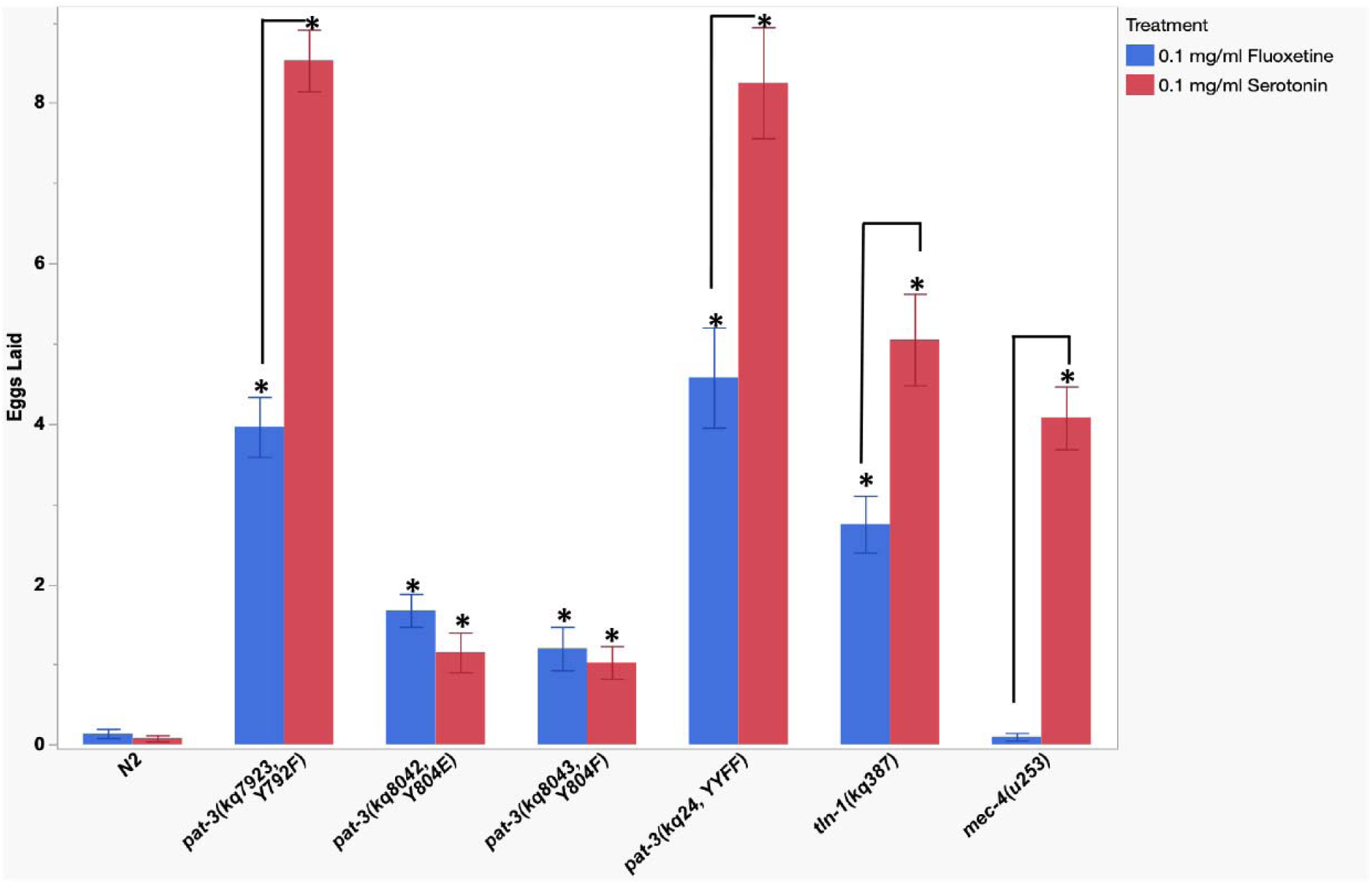
*pat-3* NPxY mutants display hypersensitivity to serotonin and fluoxetine. Egg-laying assays were repeated in *pat-3* NPxY mutants at a concentration of serotonin and fluoxetine, which does not stimulate egg laying in N2 (0.1 mg/ml serotonin or fluoxetine). Multiple adult worms (n=40) were used for each experiment. Brackets indicate significant differences between the treatments. Results significantly different from N2 were marked with * (*p* < *0.05*). *All Pairwise Comparisons - Tukey HSD.

The membrane-proximal NPxY motif is known as a binding site for talin, a molecular bridge connecting the actin cytoskeleton to integrin ^33^. Our previous study created a tryptophan (W) to alanine (A) mutation at position 387 in the F3 FERM domain of TLN-1/talin, denoted as *tln-1(kq387, W387A)*. Structural analysis of the talin-integrin interaction demonstrated that W359 of mammalian talin, homologous to W387 in *tln-1*, binds to the proximal NPxY of β3 integrin tail. However, *tln-1(kq387)* showed mild egg-laying defects or delays in motility ^34^. The *tln-1(kq387)* was treated with 0.1 mg/ml serotonin and fluoxetine for egg-laying assays (Figure 4, supplementary table 4). Each treatment showed increased egg laying during the incubation, suggesting that the focal adhesion protein TLN-1 is also linked to serotonin hypersensitivity.

### Egg retention of NPxY mutants was compared to N2

Due to the delayed egg release, Egl mutants harbor more unlaid eggs in the uterus than N2 ^35^. Fertilized eggs are released from the uterus before reaching the 26-cell stage embryo ^36^. We scored the number of eggs in the uterus (Figure 5, Supplementary Table 5) from the Day 2 egg-laying worms. The number of unlaid eggs in most NPxY mutants was comparable to N2 egg retention. However, *pat-3(Y792F)* displayed fewer unlaid eggs in the uterus than N2 (*p < 0.0019*, n = 50) or the other mutants. This analysis showed that *pat-3(Y792F)* retained fewer eggs in the uterus while laying more eggs in serotonin stimulations, suggesting that egg laying was accelerated. However, *pat-3(Y792F)* maintained egg retention (average 10 eggs per animal), which is considered within a typical range of egg bearing ^37^. Therefore, the statistical significance failed to support the argument that the *pat-3(Y792F)* mutation accelerated egg release in the uterus.

**Figure 5.**
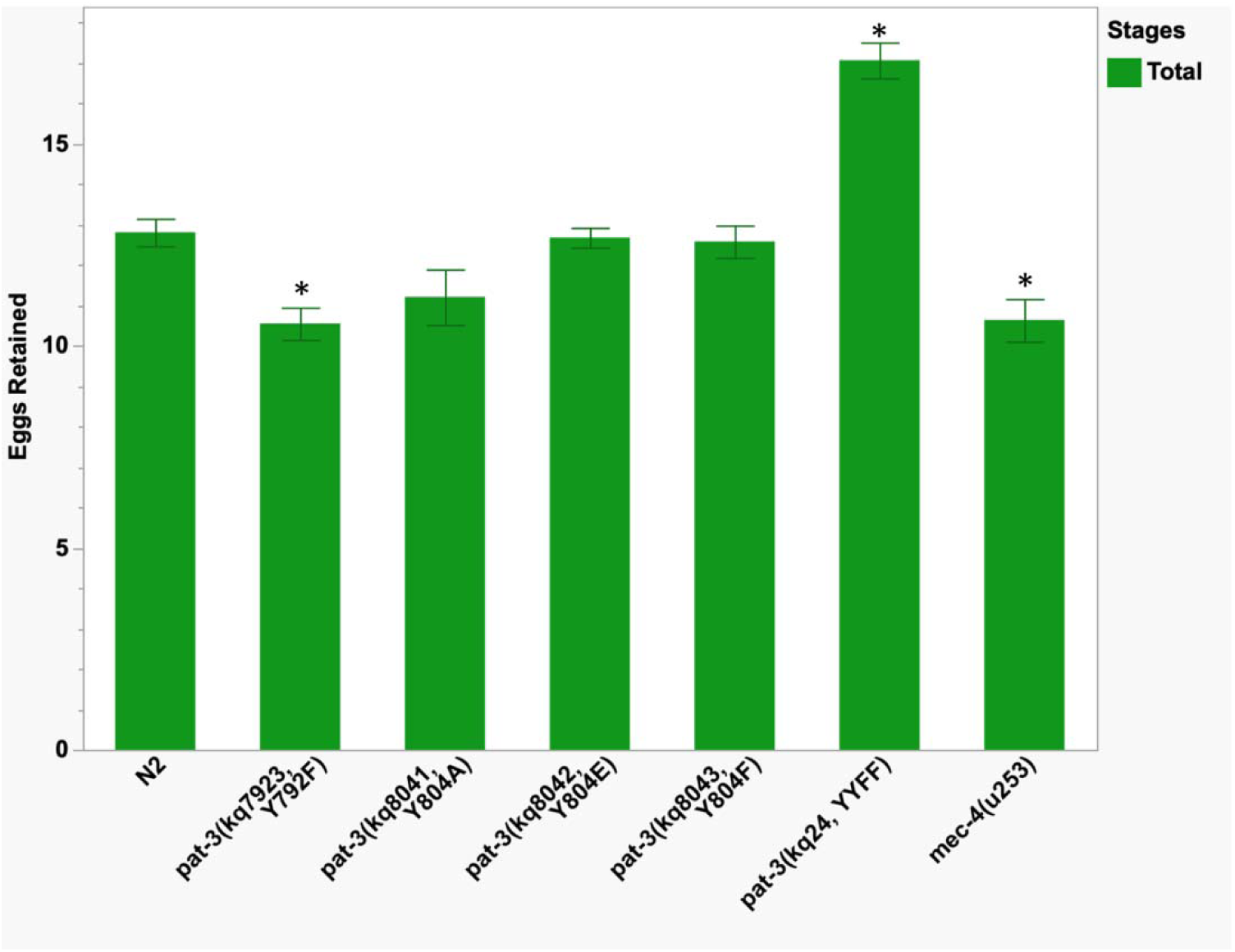
Egg retention assays of integrin mutants. Results significantly different from N2 were marked with * (*p* < 0.05). Multiple worms (n = 35 or 50) were used for each scoring. Bars represent total eggs present at the time of assay. * All Pairwise Comparisons – One-way analysis of variance.

### Double non-phosphorylatable NPxY showed a distinct Egl phenotype

In mammalian studies, the same knock-in of YY to FF in mouse β1 and β3 integrins showed no overt behavioral phenotypes ^6^. The *pat-3(kq24, YYFF)* laid a comparable number of eggs in serotonin or fluoxetine during the sixty-minute incubation, suggesting that the egg-laying appeared similar to N2 (for serotonin, *p* < 0.99, and fluoxetine, *p < 0.88*) (Figure 3, Supplementary Table 3). However, the egg retention of *pat-3(YYFF)* showed a higher number of unlaid eggs in the uterus than in N2 (*p* < 0.0001). The *pat-3(YYFF)* carried about thirty percent more unlaid eggs than N2 animals (Figure 5). This result suggested that *pat-3(YYFF)* animals also delayed egg-laying while capable of vulva contraction in response to serotonin and fluoxetine (Figure 3). The *pat-3(YYFF)* was also tested for serotonin sensitivity because it includes the proximal non-phosphorylation Y792F. Likewise, *pat-3(YYFF)* displayed a substantial egg-laying increase by the reduced serotonin and fluoxetine concentration, 0.1 mg/ml (Figure 4).

The serotonin hypersensitivity of Y792F and YYFF mutations prompted us to test the ‘distal’ non-phosphorylation, *pat-3(Y804F)*, which displayed only a slight increase (average 1.4 eggs per hour) in egg laying and failed to show the same level of hypersensitivity as *pat-3(Y792F)* or *pat-3(YYFF)* (Figure 4).

### The mechanosensation of βpat-3 integrin NPxY mutants

*C. elegans* egg laying is a complex behavior that requires the coordination of muscles and neurons, including vulva and uterine muscles and HSN and VC motor neurons ^37^. While the current study primarily focuses on contractile tissues, our results imply the potential involvement of integrin in neurons associated with egg laying.

In *pat-3(Y804E)*, serotonin treatment failed to improve egg laying to N2 levels, although egg retention fell within the normal range of 10 to 15 ^37^ (Figures 3 and 4). The *pat-3(Y792F)* and *pat-3(Y804A)* mutants were more or equally sensitive to serotonin compared to N2 (*p* < 0.0001 and *p* < 0.87, respectively), but both were less sensitive to fluoxetine than N2 (*p* < 0.0001 and *p* < 0.01). In contrast, *pat-3(YYFF)* displayed egg laying comparable to N2 in both treatments while exhibiting a higher accumulation of unlaid eggs than N2 (*p* < 0.0001). This suggests that their altered behavior may result from impaired ability to respond to serotonin or to initiate egg laying due to decreased sensitivity to stimuli from surroundings (Figure 3).

Several studies demonstrated the connections among mechanosensation, egg laying, and integrin. Chalfie *et al.* (1985) identified a pair of PLM touch neurons extending along the posterior lateral side and synapse with the HSN motor neurons ^38^. Vibrational stimulation inhibited egg laying, and this response was dependent on the ALM and PLM touch neurons ^39^. Ravie *et al.* (2018) proposed that mechanical pressure from egg accumulation in the uterus triggers egg laying through a potential homeostatic mechanism, mediated by a touch neuron ^40^. Chen and Chalfie (2014) demonstrated that *pat-3* mosaic animals exhibited mechanosensation defects involved in ALM ^22^. These possible links among mechanosensation, egg laying, and integrin led us to hypothesize that the phosphorylation of integrin NPxY is necessary for the touch response, and that mutations in PAT-3 NPxY phosphorylation would result in a defective response to mechanical stimuli ^41^.

Therefore, we evaluated the NPxY mutants using gentle touch analyses ^42^ (Figure 6, Supplementary Table 6). Well-fed adult animals were touched (gentle prodding) on the anterior or posterior half of the body ^43,44^. The scoring involved assessing backing (anterior touch) or sprinting forward movement (posterior touch) after the gentle touch stimuli (see Experimental Procedures) ^23^. Our results revealed that most NPxY mutants exhibited significantly decreased touch responses on the anterior or posterior side of the body (Figure 6). However, *pat-3(kq8041, Y804A)* and *pat-3(kq8043, Y804F)* appeared wild-type in response to gentle touch stimuli.

**Figure 6.**
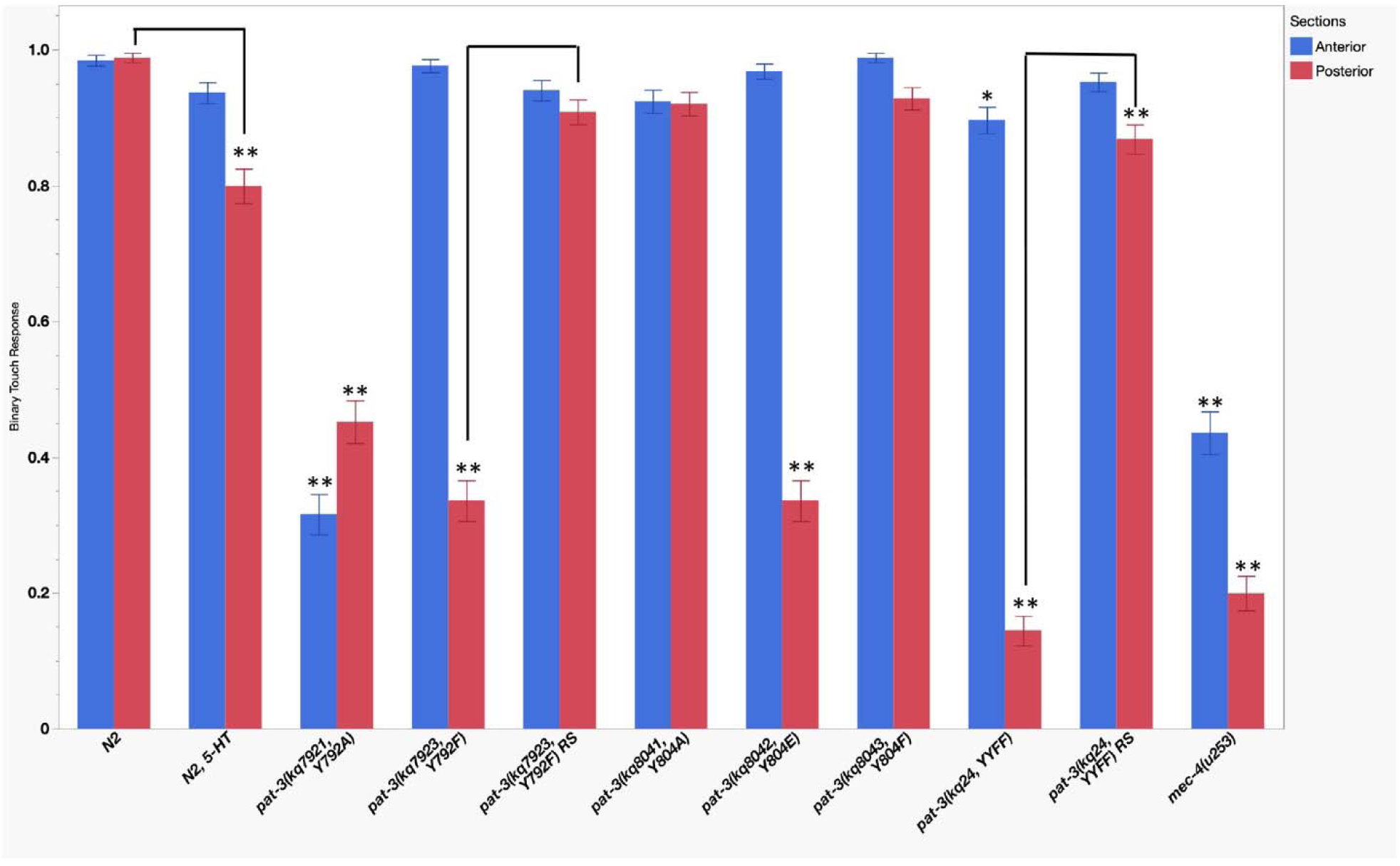
Mechanosensation analysis of *pat-3* NPxY mutants and *mec-4*. The single asterisk (*) mark on *pat-3(kq24)* anterior indicates *p* = 0.048, while all other results significantly different from N2 are marked with ** (*p* < 0.05). Brackets indicate significant differences between the treatment with serotonin (5-HT) or reserpine (RS) and no treatment. All Pairwise Comparisons – Tukey HSD analyzed the averages of each test (n = 50). 5-HT: serotonin treatment (0.5 mg/ml); RS: reserpine (60 μM). Worms were treated with the drugs for 24 hours before touch assays.

Two mutants, *pat-3(kq7923, Y792F)* and *pat-3(kq8042, Y804E),* exhibited significant insensitivity to gentle touch (Figure 6, *p < 0.0001* compared to N2 posterior), with the defect predominantly affecting the posterior ends. To strengthen the analysis of the role of membrane-proximal NPxY in mechanosensation, *pat-3(kq7921, Y792A)*, a tyrosine removal mutation, was generated (see Experimental Procedures). The *pat-3(Y792A)* mutation caused a disruptive change, abolishing the intact membrane-proximal NPxY structure. Although *pat-3(Y792A)* animals displayed severely uncoordinated movement with irregular and patchy dense bodies (Figure 2E), they were viable and fertile. As expected, the touch response in *pat-3(Y792A)* was severely defective in the anterior and posterior bodies, suggesting that the proximal NPxY motif is also essential for motility and mechanosensation (Figure 6). The non-phosphorylatable NPxY mutants, *pat-3(kq7923, Y792F)* and *pat-3(kq24, YYFF),* displayed a severe posterior touch insensitivity (*p* < 0.0001). The mechanosensation defects coincided with phenylalanine (F) or alanine (A) in the proximal motif, suggesting that phosphorylated proximal NPxY is required for mechanosensation.

### The connection between mechanosensation and egg-laying behaviors

Our current study revealed the involvement of the *pat-3* NPxY motif in egg laying, which modulates serotonin sensitivity, while identifying the role of NPxY in mechanosensation. Therefore, we examined a mechanosensation mutant for egg laying to explore the linkage between touch sensitivity and egg-laying behaviors. The *mec-4(u253)*, a null allele of the MEC-4/ENaC mechanoreceptor ^45^, demonstrated severe touch defects in both ends (Figure 6). While the egg laying of *mec-4(u253)* in serotonin was not different from the wild type (*p < 0.995*, compared to N2), it failed to respond to fluoxetine (*p < 0.0001*) (Figure 3), suggesting that *mec-4(u253)* has a lowered serotonin level or defective CeSERT/MOD-5 serotonin transporter. Therefore, the sensitivity to serotonin or fluoxetine was also tested by treating *mec-4(u253)* with the reduced concentrations of serotonin and fluoxetine (Figure 4). The hypersensitivity experiment showed *mec-4(u253)* laid eggs in 0.1 mg/ml serotonin, comparable to *mec-4* in 3.0 mg/ml. Our data suggested that the mechanosensation channel on TRN is linked to serotonin sensitivity.

Young adult N2 were grown on serotonin-containing (0.5 mg/ml) NGM plates to address the linkage between serotonin sensitivity and mechanosensation for 24 hours. The touch sensitivity of the serotonin-grown N2 showed significantly decreased touch response in the posterior body (*p* > *0.0001*) (Figure 6). In another experiment, *pat-3(kq7923, Y792F)* and *pat-3(kq24, YYFF)* were grown on NGM plates containing 60 μM reserpine, which is a VMAT (vesicular monoamine transporter) inhibitor previously known for depleting monoamine neurotransmitters, serotonin and dopamine, at the presynaptic ends ^27^. In *C. elegans*, reserpine, an inhibitor of CAT-1/VMAT, blocks dopamine and serotonin uptake into synaptic vesicles ^27^, extends its lifespan, and confers thermotolerance^46^ at a concentration of 60 μM. We expected that the reserpine treatment would reduce serotonin release in the neurons. The *pat-3(YYFF)* and *pat-3(Y792F)* mutants were grown on plates containing 60 μM reserpine for 24 hours. The reserpine treatment in the *pat-3* alleles significantly suppressed their defective mechanosensation (Figure 6), suggesting that serotonin negatively regulates the mechanosensation and that serotonin depletion rescues the NPxY mutation-induced mechanosensation defects.

## DISCUSSION

Our study reports new functions of integrin NPxY motifs. We observed varied egg-laying responses to serotonin and fluoxetine among *pat-3* alleles: *pat-3(kq8042, Y804E), pat-3(kq7923, Y792F)*, and *pat-3(kq24, YYFF)*. Functional analysis of the membrane-distal NPxY motif reveals that the phosphomimetic *pat-3(Y804E)* mutant significantly reduces egg laying under both treatments. The double non-phosphorylatable *pat-3(YYFF)* allele shows regular egg laying to serotonin and fluoxetine but retains more unlaid eggs in the uterus. In contrast, the proximal non-phosphorylatable *pat-3(Y792F)* mutant exhibits reduced fluoxetine sensitivity, increased egg laying in serotonin. Additionally, *pat-3(Y792F)* and *pat-3(YYFF)* mutants display hypersensitivity to serotonin and fluoxetine. Beyond egg-laying behavior, these NPxY mutants, including *pat-3(kq7921, Y792A)*, exhibit defective mechanosensation, dependent on serotonin levels. This suggests that timely phosphorylation of the NPxY motif is critical for regulating vulval muscle contraction and touch receptor neuron function.

### The phosphomimetic pat-3(kq8042, Y804E)

Our data demonstrate that *pat-3* NPxY is essential for responding to serotonin and fluoxetine. Our findings reveal that tyrosine phosphorylation disrupts vulva contraction, as observed in *pat-3(Y804E)*, suggesting that the membrane-distal Y^804^ residue is a critical switch for serotonin and fluoxetine responses. This result suggests that the distal NPxY^804^ potentially interacts with MOD-5/CeSERT or serotonin receptors. Phosphorylation of distal NPxY^804^ may impair the function of MOD-5 or multiple serotonin receptors, leading to impaired contraction of the vulva muscles. Notably, Carneiro *et al.* (2008) reported that αIIbβ3 platelet integrin binds to the C-terminal domain of platelet SERT, where active β3 integrin enhances the serotonin transporter activity of the SERT ^47^. Interestingly, the egg retention of the phosphomimetic *pat-3(kq8042, Y804E)* is similar to that of N2, indicating that constitutive phosphorylation of the distal motif specifically reduces vulva muscle contraction. Fluorescence microscopy (Figure 2) reveals a wild-type distribution of TLN-1 in the body wall muscle of the *pat-3(Y804E)* mutant, demonstrating that phosphorylation of the distal NPxY is dispensable for muscle organization. We conclude that an unknown factor, parallel to or downstream of *pat-3(Y804E)*, such as *goa-1/Go*α ^48^, likely influences serotonin sensitivity and muscle contraction. Segalat *et al.* show that the activated GOA-1, a Goα protein homolog expressed in muscle or HSN, inhibits egg laying ^48^. The PAT-3 distal NPxY phosphorylation potentially activates GOA-1 or other Gα subunits and, consequently, inhibits the contraction of the vulva muscle.

### *pat-3(kq24, YYFF)*, the double non-phosphorylation

The *pat-3(YYFF)* exhibits hypersensitivity to serotonin and higher uterine egg retention. This observation suggests that *pat-3(YYFF)* is defective in initiating egg laying, possibly due to impaired ability to detect mechanical pressure from the oviparous uterus. This defect could arise from an abnormal serotonin level to trigger vulva contraction ^35^ or the failure of the utero-vulva feed-forward mechanism to stimulate egg-laying via serotonin secretion ^49^. Our results on *pat-3(YYFF)* indicate that the egg retention of the YYFF mutation is a combined effect created by an interaction between Y^792^ and Y^804^. The Y792F or Y804F mutation appears normal in egg retention (Figure 4), which supports a potential interaction between the two NPxY motifs. Further studies will elucidate the molecular nature of any possible interactions between the two NPxY motifs in β-integrin.

The hypersensitivity of membrane-proximal non-phosphorylation, *pat-3(kq7923, Y792F)*. In contrast, the *pat-3(Y792F)* mutant strongly responds to serotonin, but has a reduced reaction to fluoxetine (Figure 4). In *C. elegans*, the endogenous serotonin is produced and secreted from NSM (neurosecretory motor in the head), ADF (amphid dual ciliated ending F sensory), and HSN (hermaphrodite-specific motor) neurons ^50^. There is no evidence of serotonin production in muscle cells, suggesting that the NPxY phosphorylation regulates serotonin production or secretion in non-muscle cells. Our results also indicate the importance of the membrane-proximal NPxY in serotonin sensitivity. The hypersensitivity to serotonin in *pat-3(Y792F)* and *pat-3(YYFF)* suggests that Y^792^ regulates the sensitivity to serotonin. Walser *et al.* (2017) report that density-dependent phosphatase (*dep-1*) unphosphorylates the membrane-proximal NPxY. Our finding suggests that the proximal NPxY is kept unphosphorylated during active egg laying.

### Serotonin hypersensitivity shows that integrin regulates egg laying in a serotonin-dependent or independent manner

The molecular nature of hypersensitivity to serotonin and fluoxetine is unknown and requires further studies. However, there are several possible explanations. Our search finds that the serotonin hypersensitivity is similar to the *mod-5/CeSERT* loss-of-function phenotypes. Ranganathan *et al.* show the serotonin-specific hypersensitivity of *mod-5* mutants in hyper-enhanced slowing behavior ^31^. In contrast, fluoxetine stimulation of egg laying in *tph-1(gm280)* mutants indicates that egg laying could be induced without the endogenous serotonin. Our analysis suggests that *pat-3(Y792F)* and *pat-3(YYFF)* are serotonin hypersensitive, indicating that the proximal NPxY-dependent egg laying can be serotonin-independent. Kullyev et al. (2010) describe that fluoxetine binds to *ser-5* and *ser-7* receptors and causes serotonin-independent behaviors. We speculate that fluoxetine independently stimulates serotonin receptors on muscles, which are enhanced by Y792F and YYFF mutations. However, our data indicate that *pat-3(Y792F)* and *pat-3(YYFF)* showed less sensitivity to fluoxetine than serotonin. This result suggests an alternative scenario that the level of serotonin might be increased in *pat-3(Y792F)* and *pat-3(YYFF)* due to potential inhibitory effects of the NPxY mutations in mod-5/CeSERT functions.

### Several NPxY mutants display defective mechanosensation

The *pat-3(kq7921, Y792A)* exhibits mechanosensation defects in both the anterior and posterior regions, whereas *pat-3(kq7923, Y792F)*, *pat-3(kq8042, Y804E)*, and *pat-3(kq24, YYFF)* show the defect primarily in the posterior half. Our analysis suggests that NPxY phosphorylation modulates mechanosensation (see Figure 5). While Chen *et al.* (2014) demonstrated that *pat-3(-)* mosaic in ALM neurons exhibits touch insensitivity ^22^, many NPxY mutants exhibit posterior touch defects, suggesting that the role of the NPxY motif is biased to the PLM neurons. This aligns with the βpat-3 localization of ALM and PLM neurons in Gettner *et al.* ^9^ and Amezquita *et al.* ^51^.

Overall, our data suggest that touch stimuli induce phosphorylation of the NPxY motifs. The touch insensitivity at the posterior body in *pat-3(YYFF)* and *pat-3(Y792F)* indicates that phosphorylation of NPxY motifs may contribute to the mechanosensation. Mechanical stimuli might simultaneously or sequentially phosphorylate each NPxY, transmitting mechanical information to mechanotransduction targets within the cell. The NPVY^804^ motif appears to bind a downstream molecule, UNC-112/kindlin. The phosphorylated NPVY^804^ may fail the interaction between UNC-112/kindlin and the membrane distal NPxY ^52^. We propose that the *pat-3(Y804E)* mechanosensation defect arises from impaired interaction with UNC-112/kindlin.

Likewise, phosphorylation of the proximal NPIY^792^ inhibits binding to TLN-1/talin ^53^. In our study, however, the mechanosensation of *pat-3(Y792F)* is defective in the posterior end, suggesting phosphorylation of proximal NPIY^792^ transiently occurs in PLM and affects its ability to sense mechanical stimuli. In contrast, tyrosine removal or single non-phosphorylatable mutations in the distal motif, such as *pat-3(Y804A)* and *pat-3(Y804F)*, show wild-type touch sensitivity in the anterior or posterior end. This suggests that these mutations have minimal impact on the involvement of distal NPxY in mechanosensation.

How does βPAT-3 integrin contribute to mechanosensation? One straightforward interpretation is that the NPxY mutants cause unknown muscle anomalies, thereby reducing the motor activity of muscles in response to touch stimuli. However, the βPAT-3 protein is expressed in all tissues, particularly muscles and neurons ^9^. Therefore, one cannot rule out the potential role of βPAT-3 NPxY in neurons. Previous analysis shows that the *pat-3* integrin directs the posterior-to-anterior migration of PLM axons during development ^54^. This suggests that the mechanosensation defect observed in NPxY mutants stems from impaired PLM axon outgrowth. Consequently, stranded PLM axons fail to detect touch stimuli in the posterior body. Another possibility is that integrin may act as a direct mediator of mechanotransduction, potentially interacting with MEC-4 or other touch receptors ^55^. Tyrosine phosphorylation of the proximal NPIY^792^ transiently occurs, facilitating interaction with MEC-4. Upon touch stimulation, the MEC-4 channel may induce phosphorylation of the proximal motif, subsequently triggering the release of neurotransmitters at the PLM synapse to inhibit motor neurons such as HSN ^42,49^.

Our analysis shows that exogenous serotonin reduces the touch sensitivity of N2, while inhibition of neurotransmitter secretion rescues the mechanosensation of NPxY mutants. This demonstrates that serotonin reduces sensory response while promoting egg laying. This result suggests that integrin NPxY phosphorylation potentially orchestrates the secretion and production of serotonin. Integrin maintains serotonin at a level, presumably, in neurons. This means that the phosphorylation or dephosphorylation of *pat-3* NPxY modulates serotonin levels in the tissues. Thus, non-phosphorylation of the proximal NPxY causes a general increase in serotonin, which attenuates mechanosensation.

In mammalian studies, β3 integrin YYFF knock-in mice exhibited a mild defect in clot retraction and a tendency to rebleed ^8^. However, other studies with β1 YYFF knock-in mice failed to detect any developmental defects in the mutant animals ^6^. Our study presents hypersensitive egg laying and defective mechanosensation phenotypes associated with the Y792F, Y804E, and YYFF mutations. This suggests that the membrane-proximal and distal NPxY motifs in mice β1 or β3 integrin play a role in reproductive and sensory behaviors similar to those of βPAT-3 *C. elegans* integrin.

## EXPERIMENTAL PROCEDURES

### Animals and Genetics

All animals were fed with *E. coli*, OP50, and cultured at room temperature (21°C to 23°C) at all times. *C. elegans* N2 worm, AH3437 GFP::*tln-1*, and *mec-4(u253),* were obtained from Caenorhabditis Genetics Center at St. Paul, MN. The CRISPR-Cas9 edited worms*, pat-3(kq8041, Y804A), pat-3(kq8042, Y804E),* and *pat-3(kq8043, Y804F)* were previously generated and described ^12,56^. All *pat-3* NPxY mutants and other lines are listed in Supplementary Table 1 (Table S1).

Genetic double mutant lines (Table S1) were generated by crossing homozygote males from AH3437 GFP::*tln-1* ^28^ to each NPxY edited line hermaphrodite. From the F2 generation, a worm with 100% green (F3) progeny and the *pat-3* NPxY allele homozygosity (by PCR genotyping) was selected for further studies.

### Phenotype Analysis and Fluorescence Imaging

Fecundity analyses of N2, *pat-3 NPxY*, and other mutants were conducted as follows. Eggs were collected from worms grown on NGM plates (Day -2). After 48 hours (Day 0), the hatched larvae were transferred to individual NGM plates, with one worm per plate. At 72 hours (Day 1), each worm was transferred daily to a new NGM plate for four additional days. After the worm was removed and placed onto a new plate, the progeny on the previous plate were allowed to grow for one more day. The viable progeny were then stored at 4°C, and the number of progeny on each plate was subsequently counted and recorded for analysis.

For egg-laying assays, an adult worm was placed in 100 μl of M9 buffer per well in a 96-well microtiter plate containing fluoxetine (0.5 mg/ml, Sigma-Aldrich) or serotonin (3 mg/ml). After a 15-minute acclimation period, the number of eggs laid was counted 60 minutes later. For hypersensitivity egg-laying assays, adult worms were placed in M9 buffer containing 0.1 mg/ml fluoxetine or serotonin. For egg retention assays, eggs were transferred to an OP50-seeded plate and grown for four days at room temperature.

Adult worms were then placed on a microscope slide containing 50 to 100 μl of M9 buffer, and eggs were extruded using a 26-gauge needle. The extruded eggs were counted and categorized by developmental stage, with eggs at the bean-shaped or later embryonic stages scored as late-stage embryos.

For mechanosensation assays, following the protocol by Hobert *et al.* (1997), young adult animals that had been fed *E. coli* OP50 for two generations without starvation were gently stroked with eyebrow hairs ten times—five times on the anterior body and five times on the posterior body ^23^. Responses such as backing movement to anterior touch and sprinting or twisting to posterior touch were scored as 1, while no responses to touch were scored as 0. For serotonin mechanosensation assays, worms were grown on NGM plates containing 0.5 mg/ml serotonin or 60 μM reserpine (Sigma-Aldrich) for 24 hours before the touch assays. Sample images were taken using a Nikon Eclipse Ni-U fluorescence microscope and a Coolsnap Dyno digital monochrome camera (Photomerics, Tucson, AZ). Photograph images were analyzed using NIS elements (Nikon, Melville, NY).

### The generation of CRISPR-Cas9 knock-in C. elegans

The CRISPR-Cas9 system was used to edit the *pat-3* locus to create the *kq24*, YY792/804FF (YYFF) mutation, and other alleles. For *kq24*, the two crRNA, ZQPATY1(CGAGAACCCAATCTACAAAC) and ZQPAT3B (TTTAAAAATCCAGTATACGC) were obtained from the exon 8 of the *pat-3* gene covering the membrane-proximal and distal NPxY and purchased from IDT Inc., Coralville, IA. The homologous directed repair template (taaatttatcaaattatcatttttcagAACGAGAATCCAATTTTTAAGCAAGCCACGACAACATTCAAGAACCCGGTTTTTGCAGGAAAAGCCAACTAAatagtttttatccttatatt) spans 48 bases upstream and 38 bases downstream of the target site, tyrosine (Y^792^ and Y^804^).

Mutations modified the third position of codons in the crRNA target sequence to a synonymous codon. The repair DNA templates, crRNA, tracrRNA, and Cas9 nuclease were custom-made by IDT Inc., Coralville, IA. A mixture of template DNA, two crRNA (ΖQAPTY1 and ZQPAT3B), tracrRNA (cat. no. 1073190), and Cas9 protein (cat. no. 1081058) was annealed at room temperature. The mixture was micro-injected into the syncytial gonad of N2 animals (P0) with *dpy-10* co-CRISPR ^57,58^. Homozygous mutants were crossed back to N2 two times. The mutation was confirmed by sequencing DNA molecules extracted from homozygous animals ^12^. The generation of other edits was carried out similarly to the techniques mentioned above. The nucleotide sequences of crRNA and templates are listed in Supplementary Table S2 (Table S2).

### Statistical Analysis

Statistical analysis of multiple *C. elegans* strains was conducted using All Pairwise Comparisons - Tukey HSD (Honestly Significant Difference) method ^59^. The egg-laying assay utilized a Poisson regression model, while the regular and late-stage egg retention assays employed the Comparison of Means. A significance level was set at *p < 0.05.* Touch sensing assays were subjected to logistic regression and all pairwise comparisons for Tukey HSD. All data were analyzed using JMP Pro (version 17.0, SAS Institute, Cary, NC) software. Pairwise comparison data with *p > 0.05* can be found in Supplementary Tables 3, 4, 5, and 6.

## Acknowledgments

The Office of Engaged Learning at Baylor University URSA Research Fund supported this work. The *pat-3* mutants were generated for the BIO4108 Laboratory Class. The authors acknowledge Catherine Ravikumar, Arlyn Alcid, and Jeancarlo Gutierrez for their help during the initial stage of the project. The authors also appreciate Drs. Erin Cram and Ilya Ruvinsky for their comments on the initial draft.

