## supplementary figures and tables 1 & 2 for "The β integrin modulates serotonin sensitivity via NPxY motifs to regulate egg laying and mechanosensation behaviors in *C. elegans*"

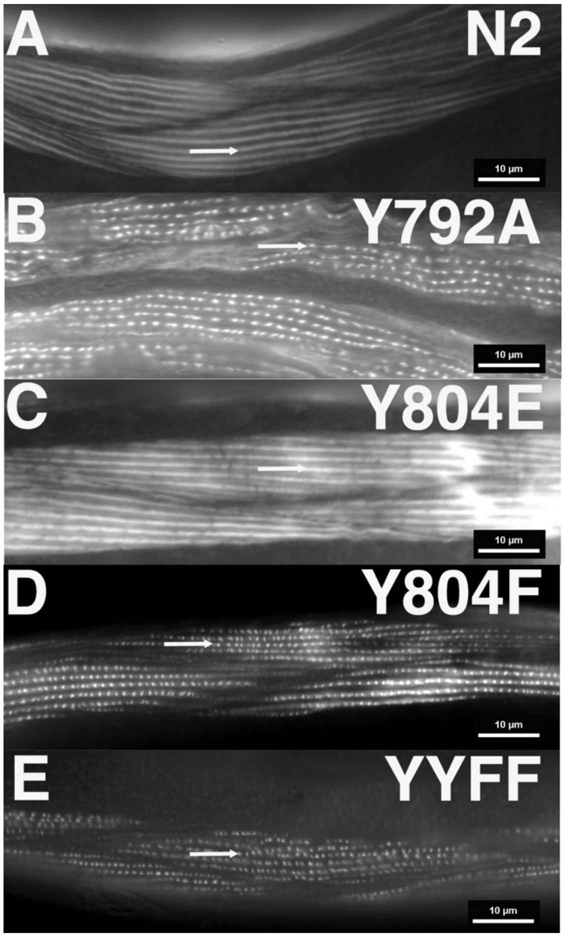


Figure S1. Rhodamine-conjugated phalloidin staining of NPxY mutants.

The four *pat-3* NPxY mutants were stained with rhodamine-conjugated phalloidin. Prior to the staining, worms were fixed with acetone and methanol. In every phots, horizontal dots or lines represent actin filaments (arrows). Except for panel B, all other images show regularly striated actin filament patterns. Although panel B shows filament patterns, it displays a few interruptions in staining and distortion of filaments. All images are taken under a 60X objective lens. Homozygous mutants are shown here. A. body wall muscle of N2 adult worm (control), B. *pat-3(kq7921, Y792A),* C. *pat-3(kq8042, Y804E),* D. *pat-3(kq8043, Y804F),* E. *pat-3(kq24, YYFF),* Bars = 10 µm.

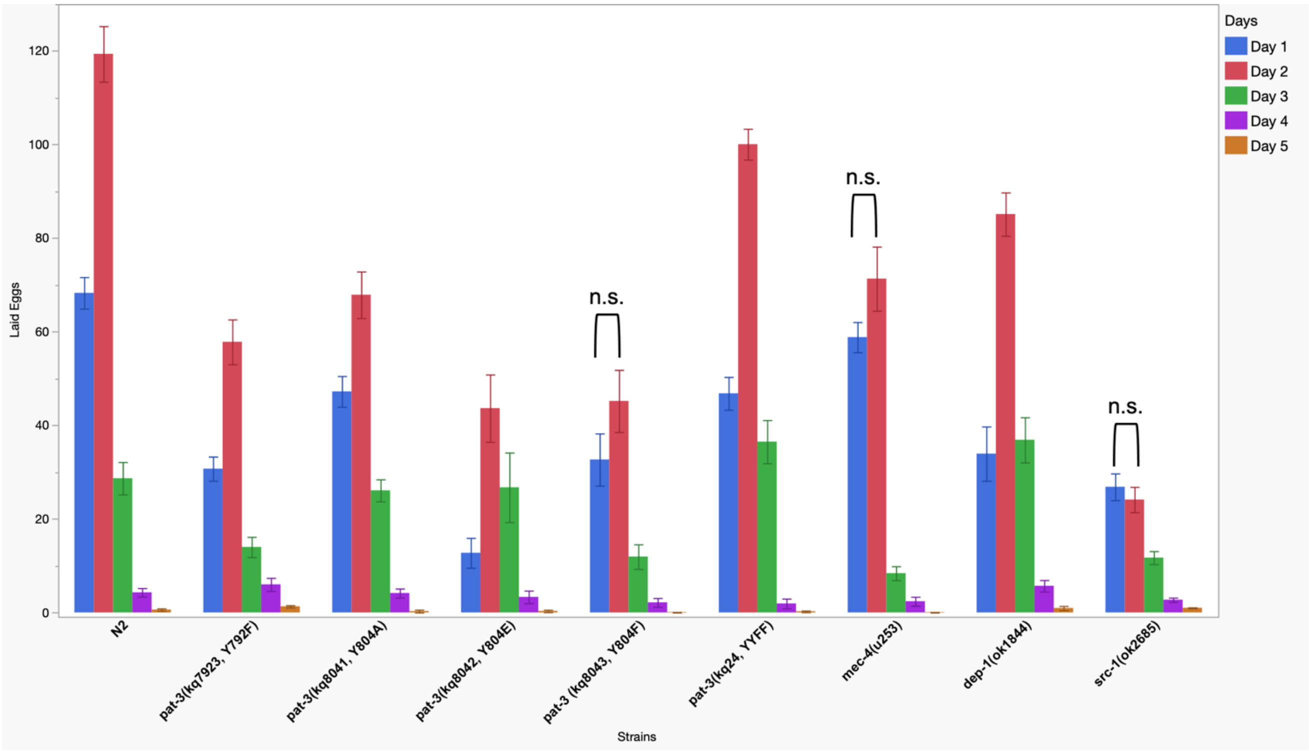


Figure S2. Fecundity analysis of *pat-3* NPxY mutants, *mec-4*, *dep-1*, and *src-1*.

The number of eggs laid daily was compared in each strain (13 to 22 worms were used in each strain). N.S. indicates $p > 0.05$ in ANOVA with All Pairwise Comparisons – Tukey HSD.

**Supplementary Table 1. CRISPR edits and mutant strains used for this study**

| | |
|---|---|
| BU7921 | *pat-3(kq7921,Y792A),* from this study |
| BU7923 | *pat-3(kq7923,Y792F),* from this study |
| BU8041 | *pat-3(kq8041,Y804A),* available in CGC |
| BU8042 | *pat-3(kq8042,Y804E),* available in CGC |
| BU8043 | *pat-3(kq8043,Y804F),* available in CGC |
| BU24 | *pat-3(kq24,YYFF)*, from this study |
| BU30 | *GFP::tln-1;pat-3(kq7921,Y792A)*, from this study |
| BU28 | *GFP::tln-1;pat-3(kq8041,Y804A)*, from this study |
| BU25 | *GFP::tln-1;pat-3(kq8042,Y804E)*, from this study |
| BU26 | *GFP::tln-1;pat-3(kq24,YYFF)*, from this study |
| TU253 | *mec-4(u253)* from CGC |

**Supplementary Table 2. The *pat-3* Exon8 Y to A, E, or F CRISPR-CAS9 reagents**

crRNA and repair template used to generate pat-3 edits. Red indicates the mutations.

| ssDNA Oligo name | Sequence (5'-3') |
|---|---|
| ZQPATY1 (crRNA) | CGAGAACCCAATCTACAAAC |
| ZQPAT3B (crRNA) | TTTAAAAATCCAGTATACGC |
| Repair template Y1F | taaatttatcaaattatcatttttcagAACGAGAATCCAATTTTTAAGCAAGCCACGACAACATTTAAAAATCCAGTATACGCTGG |
| Repair template Y1A | taaatttatcaaattatcatttttcagAACGAGAATCCAATTGCTAAGCAAGCCACGACAACATTTAAAAATCCAGTATACGCTGG |
| Repair template YYFF | taaatttatcaaattatcatttttcagAACGAGAATCCAATTTTTAAGCAAGCCACGACAACATTCAAGAACCCGGTTTTTGCAGGAAAAGCCAACTAAatagtttttatccttatatt |

PCR primers for genotyping.

| PCR primer name | Sequence (5'-3') |
|---|---|
| PCR-WT-R-Y1 | GCCTGTTTGTAGATTGGG |
| PCR-R-Y1F | GTGGCTTGCTTAAAAATTGGA |
| PCR-R-Y1A | GGCTTGCTTAGCAATTGGA |
| PCR-R-Y2F | GCAAAAACCGGGTTCTTG |
| PCR-R-Y2E | CTTCAACCGGGTTCTTG |
| PCR-R-Y2A | CAGCAACCGGGTTCTTG |
| PAT3MCRF | CATGATAGATCCGAATACGC |
| PAT3R3UTR | acaatttatcgctaaatactcgtt |
| PAT3TTWT | AGCGTATACTGGATTTTTAAATGTTGTC |
