## Supplementary Table 3 for "The β integrin modulates serotonin sensitivity via NPxY motifs to regulate egg laying and mechanosensation behaviors in *C. elegans*"

**Egg laying** new **p>0.05**

| Strains | Treatment | -Strains | -Treatment | Difference | Std Error | t Ratio | Prob>\|t\| | Lower 95% | Upper 95% |
|---|---|---|---|---|---|---|---|---|---|
| N2 | Fluoxetine | pat-3(kq8043, Y804F) | Fluoxetine | 0.22832 | 0.0719818 | 3.17 | 0.0889 | -0.01404 | 0.47069 |
| pat-3(kq8043, Y804F) | Fluoxetine | pat-3(kq8043, Y804F) | Serotonin | -0.21637 | 0.0684796 | -3.16 | 0.092 | -0.44694 | 0.0142 |
| pat-3(kq8043, Y804F) | Fluoxetine | pat-3(kq7923, Y792F) | Fluoxetine | 0.23595 | 0.0771564 | 3.06 | 0.1212 | -0.02384 | 0.49573 |
| pat-3(kq7923, Y792F) | Fluoxetine | mec-4(u253) | Serotonin | -0.22024 | 0.0774224 | -2.84 | 0.2053 | -0.48092 | 0.04044 |
| pat-3(kq8041, Y804A) | Serotonin | pat-3(kq24, YYFF) | Fluoxetine | -0.18962 | 0.074754 | -2.54 | 0.3844 | -0.44132 | 0.06208 |
| pat-3(kq24, YYFF) | Fluoxetine | pat-3(kq24, YYFF) | Serotonin | 0.1972 | 0.0793796 | 2.48 | 0.4206 | -0.07007 | 0.46448 |
| pat-3(kq8041, Y804A) | Fluoxetine | pat-3(kq24, YYFF) | Fluoxetine | -0.18414 | 0.0746535 | -2.47 | 0.4332 | -0.4355 | 0.06722 |
| pat-3(kq8041, Y804A) | Fluoxetine | pat-3(kq7923, Y792F) | Fluoxetine | 0.18534 | 0.0780253 | 2.38 | 0.4995 | -0.07737 | 0.44805 |
| pat-3(kq8041, Y804A) | Serotonin | pat-3(kq7923, Y792F) | Fluoxetine | 0.17986 | 0.0781214 | 2.3 | 0.5539 | -0.08318 | 0.44289 |
| pat-3(kq24, YYFF) | Serotonin | pat-3(kq7923, Y792F) | Fluoxetine | 0.17227 | 0.0825585 | 2.09 | 0.7104 | -0.1057 | 0.45025 |
| pat-3(kq24, YYFF) | Fluoxetine | mec-4(u253) | Serotonin | 0.14924 | 0.0740232 | 2.02 | 0.7572 | -0.1 | 0.39847 |
| N2 | Fluoxetine | N2 | Serotonin | 0.14728 | 0.074686 | 1.97 | 0.7847 | -0.10418 | 0.39875 |
| N2 | Serotonin | pat-3(kq8043, Y804F) | Serotonin | -0.13533 | 0.0713166 | -1.9 | 0.8274 | -0.37545 | 0.1048 |
| pat-3(kq8043, Y804F) | Fluoxetine | pat-3(kq24, YYFF) | Fluoxetine | -0.13353 | 0.0737449 | -1.81 | 0.8709 | -0.38183 | 0.11477 |
| N2 | Serotonin | pat-3(kq8041, Y804A) | Serotonin | 0.13713 | 0.0757711 | 1.81 | 0.8713 | -0.11799 | 0.39225 |
| N2 | Serotonin | pat-3(kq24, YYFF) | Serotonin | 0.14471 | 0.0803382 | 1.8 | 0.8752 | -0.12578 | 0.41521 |

| | | | | | | | | | |
|---|---|---|---|---|---|---|---|---|---|
| N2 | Serotonin | pat-3(kq8041, Y804A) | Fluoxetine | 0.13165 | 0.075672 | 1.74 | 0.9009 | -0.12314 | 0.38644 |
| N2 | Serotonin | mec-4(u253) | Serotonin | 0.09675 | 0.0750502 | 1.29 | 0.9913 | -0.15595 | 0.34944 |
| N2 | Fluoxetine | pat-3(kq24, YYFF) | Fluoxetine | 0.09479 | 0.0736538 | 1.29 | 0.9914 | -0.1532 | 0.34279 |
| pat-3(kq8043, Y804F) | Serotonin | pat-3(kq24, YYFF) | Fluoxetine | 0.08284 | 0.070235 | 1.18 | 0.9963 | -0.15365 | 0.31932 |
| N2 | Serotonin | pat-3(kq8043, Y804F) | Fluoxetine | 0.08104 | 0.0747758 | 1.08 | 0.9984 | -0.17073 | 0.33281 |
| pat-3(kq8043, Y804F) | Fluoxetine | pat-3(kq24, YYFF) | Serotonin | 0.06367 | 0.0778307 | 0.82 | 0.9999 | -0.19838 | 0.32573 |
| N2 | Fluoxetine | pat-3(kq8043, Y804F) | Serotonin | 0.01196 | 0.0683815 | 0.17 | 1 | -0.21828 | 0.2422 |
| N2 | Serotonin | pat-3(kq24, YYFF) | Fluoxetine | -0.05249 | 0.0763867 | -0.69 | 1 | -0.30968 | 0.2047 |
| pat-3(kq8041, Y804A) | Fluoxetine | pat-3(kq8041, Y804A) | Serotonin | 0.00548 | 0.0740236 | 0.07 | 1 | -0.24376 | 0.25472 |
| pat-3(kq8041, Y804A) | Fluoxetine | pat-3(kq8043, Y804F) | Fluoxetine | -0.05061 | 0.0730044 | -0.69 | 1 | -0.29642 | 0.1952 |
| pat-3(kq8041, Y804A) | Fluoxetine | pat-3(kq24, YYFF) | Serotonin | 0.01306 | 0.0786921 | 0.17 | 1 | -0.25189 | 0.27802 |
| pat-3(kq8041, Y804A) | Fluoxetine | mec-4(u253) | Serotonin | -0.0349 | 0.0732855 | -0.48 | 1 | -0.28166 | 0.21185 |
| pat-3(kq8041, Y804A) | Serotonin | pat-3(kq8043, Y804F) | Fluoxetine | -0.05609 | 0.0731071 | -0.77 | 1 | -0.30224 | 0.19006 |
| pat-3(kq8041, Y804A) | Serotonin | pat-3(kq24, YYFF) | Serotonin | 0.00758 | 0.0787874 | 0.1 | 1 | -0.25769 | 0.27286 |
| pat-3(kq8041, Y804A) | Serotonin | mec-4(u253) | Serotonin | -0.04038 | 0.0733878 | -0.55 | 1 | -0.28748 | 0.20671 |
| pat-3(kq8043, Y804F) | Fluoxetine | mec-4(u253) | Serotonin | 0.01571 | 0.0723597 | 0.22 | 1 | -0.22793 | 0.25934 |
| pat-3(kq24, YYFF) | Serotonin | mec-4(u253) | Serotonin | -0.04797 | 0.0780944 | -0.61 | 1 | -0.31091 | 0.21498 |
| pat-3(kq8042, Y804E) | Fluoxetine | pat-3(kq8042, Y804E) | Serotonin | -0.05398 | 0.1215434 | -0.44 | 1 | -0.46322 | 0.35526 |
