## Supplementary Table 4 for "The β integrin modulates serotonin sensitivity via NPxY motifs to regulate egg laying and mechanosensation behaviors in *C. elegans*"

| Strains | Treatment | -Strains | -Treatment | Difference | Std Error | t Ratio | Prob>\|t\| | Lower 95% | Upper 95% |
|---|---|---|---|---|---|---|---|---|---|
| pat-3(kq8042, Y804E) | 0.1 mg/ml Fluoxetine | tln-1(kq387) | 0.1 mg/ml Fluoxetine | -0.49579 | 0.1549719 | -3.2 | 0.0824 | -1.0177 | 0.02612 |
| tln-1(kq387) | 0.1 mg/ml Fluoxetine | mec-4(u253) | 0.1 mg/ml Serotonin | -0.39327 | 0.1233932 | -3.19 | 0.0853 | -0.80883 | 0.02229 |
| pat-3(kq7923, Y792F) | 0.1 mg/ml Fluoxetine | tln-1(kq387) | 0.1 mg/ml Fluoxetine | 0.36464 | 0.1189177 | 3.07 | 0.1188 | -0.03584 | 0.76513 |
| pat-3(kq8042, Y804E) | 0.1 mg/ml Fluoxetine | pat-3(kq8043, Y804F) | 0.1 mg/ml Serotonin | 0.49112 | 0.1982817 | 2.48 | 0.426 | -0.17665 | 1.15889 |
| pat-3(kq7923, Y792F) | 0.1 mg/ml Fluoxetine | tln-1(kq387) | 0.1 mg/ml Serotonin | -0.24314 | 0.100005 | -2.43 | 0.4586 | -0.57994 | 0.09365 |
| tln-1(kq387) | 0.1 mg/ml Serotonin | mec-4(u253) | 0.1 mg/ml Serotonin | 0.21452 | 0.1052875 | 2.04 | 0.7434 | -0.14007 | 0.5691 |
| pat-3(kq8042, Y804E) | 0.1 mg/ml Fluoxetine | pat-3(kq8042, Y804E) | 0.1 mg/ml Serotonin | 0.37605 | 0.1914798 | 1.96 | 0.7895 | -0.26881 | 1.02091 |
| pat-3(kq8042, Y804E) | 0.1 mg/ml Fluoxetine | pat-3(kq8043, Y804F) | 0.1 mg/ml Fluoxetine | 0.33349 | 0.1890997 | 1.76 | 0.8914 | -0.30335 | 0.97034 |
| pat-3(kq7923, Y792F) | 0.1 mg/ml Fluoxetine | pat-3(kq24, YYFF) | 0.1 mg/ml Fluoxetine | -0.14436 | 0.1025426 | -1.41 | 0.9809 | -0.4897 | 0.20098 |
| pat-3(kq24, YYFF) | 0.1 mg/ml Fluoxetine | mec-4(u253) | 0.1 mg/ml Serotonin | 0.11574 | 0.1077007 | 1.07 | 0.9985 | -0.24698 | 0.47845 |
| pat-3(kq24, YYFF) | 0.1 mg/ml Fluoxetine | tln-1(kq387) | 0.1 mg/ml Serotonin | -0.09878 | 0.1020538 | -0.97 | 0.9995 | -0.44248 | 0.24491 |
| N2 | 0.1 mg/ml Fluoxetine | N2 | 0.1 mg/ml Serotonin | 0.55966 | 0.6267193 | 0.89 | 0.9998 | -1.55099 | 2.6703 |
| N2 | 0.1 mg/ml Fluoxetine | mec-4(u253) | 0.1 mg/ml Fluoxetine | 0.33711 | 0.6268389 | 0.54 | 1 | -1.77394 | 2.44816 |
| N2 | 0.1 mg/ml Serotonin | mec-4(u253) | 0.1 mg/ml Fluoxetine | -0.22254 | 0.7071459 | -0.31 | 1 | -2.60405 | 2.15896 |
| pat-3(kq7923, Y792F) | 0.1 mg/ml Fluoxetine | mec-4(u253) | 0.1 mg/ml Serotonin | -0.02863 | 0.1057614 | -0.27 | 1 | -0.38481 | 0.32755 |
| pat-3(kq7923, Y792F) | 0.1 mg/ml Serotonin | pat-3(kq24, YYFF) | 0.1 mg/ml Serotonin | 0.03306 | 0.0718505 | 0.46 | 1 | -0.20892 | 0.27503 |
| pat-3(kq8042, Y804E) | 0.1 mg/ml Serotonin | pat-3(kq8043, Y804F) | 0.1 mg/ml Fluoxetine | -0.04256 | 0.206331 | -0.21 | 1 | -0.73744 | 0.65232 |
| pat-3(kq8042, Y804E) | 0.1 mg/ml Serotonin | pat-3(kq8043, Y804F) | 0.1 mg/ml Serotonin | 0.11507 | 0.2147775 | 0.54 | 1 | -0.60825 | 0.83839 |
| pat-3(kq8043, Y804F) | 0.1 mg/ml Fluoxetine | pat-3(kq8043, Y804F) | 0.1 mg/ml Serotonin | 0.15763 | 0.2126584 | 0.74 | 1 | -0.55856 | 0.87381 |
| mec-4(u253) | 0.1 mg/ml Fluoxetine | mec-4(u253) | 0.1 mg/ml Serotonin | -3.70787 | 0.5061992 | -7.32 | <.0001* | -5.41263 | -2.0031 |
| N2 | 0.1 mg/ml Fluoxetine | pat-3(kq7923, Y792F) | 0.1 mg/ml Fluoxetine | -3.34213 | 0.384545 | -8.69 | <.0001* | -4.63719 | -2.04707 |
| N2 | 0.1 mg/ml Fluoxetine | pat-3(kq7923, Y792F) | 0.1 mg/ml Serotonin | -4.10915 | 0.3807316 | -10.79 | <.0001* | -5.39137 | -2.82693 |
| N2 | 0.1 mg/ml Fluoxetine | pat-3(kq8042, Y804E) | 0.1 mg/ml Fluoxetine | -2.4817 | 0.3971772 | -6.25 | <.0001* | -3.8193 | -1.14409 |
| N2 | 0.1 mg/ml Fluoxetine | pat-3(kq8042, Y804E) | 0.1 mg/ml Serotonin | -2.10565 | 0.4056642 | -5.19 | <.0001* | -3.47183 | -0.73946 |
| N2 | 0.1 mg/ml Fluoxetine | pat-3(kq8043, Y804F) | 0.1 mg/ml Fluoxetine | -2.1482 | 0.4045463 | -5.31 | <.0001* | -3.51062 | -0.78578 |
| N2 | 0.1 mg/ml Fluoxetine | pat-3(kq24, YYFF) | 0.1 mg/ml Fluoxetine | -3.48649 | 0.3850829 | -9.05 | <.0001* | -4.78336 | -2.18962 |

| | | | | | | | | | |
|---|---|---|---|---|---|---|---|---|---|
| N2 | 0.1 mg/ml Fluoxetine | pat-3(kq24, YYFF) | 0.1 mg/ml Serotonin | -4.0761 | 0.3819092 | -10.67 | <.0001* | -5.36228 | -2.78991 |
| N2 | 0.1 mg/ml Fluoxetine | tln-1(kq387) | 0.1 mg/ml Fluoxetine | -2.97748 | 0.3897631 | -7.64 | <.0001* | -4.29012 | -1.66485 |
| N2 | 0.1 mg/ml Fluoxetine | tln-1(kq387) | 0.1 mg/ml Serotonin | -3.58527 | 0.3844149 | -9.33 | <.0001* | -4.87989 | -2.29065 |
| N2 | 0.1 mg/ml Fluoxetine | mec-4(u253) | 0.1 mg/ml Serotonin | -3.37075 | 0.3859525 | -8.73 | <.0001* | -4.67055 | -2.07095 |
| N2 | 0.1 mg/ml Serotonin | pat-3(kq7923, Y792F) | 0.1 mg/ml Fluoxetine | -3.90178 | 0.5049784 | -7.73 | <.0001* | -5.60244 | -2.20113 |
| N2 | 0.1 mg/ml Serotonin | pat-3(kq7923, Y792F) | 0.1 mg/ml Serotonin | -4.66881 | 0.5020806 | -9.3 | <.0001* | -6.3597 | -2.97792 |
| N2 | 0.1 mg/ml Serotonin | pat-3(kq8042, Y804E) | 0.1 mg/ml Fluoxetine | -3.04135 | 0.5146631 | -5.91 | <.0001* | -4.77462 | -1.30808 |
| N2 | 0.1 mg/ml Serotonin | pat-3(kq8042, Y804E) | 0.1 mg/ml Serotonin | -2.6653 | 0.5212407 | -5.11 | <.0001* | -4.42072 | -0.90988 |
| N2 | 0.1 mg/ml Serotonin | pat-3(kq8043, Y804F) | 0.1 mg/ml Fluoxetine | -2.70786 | 0.5203711 | -5.2 | <.0001* | -4.46035 | -0.95537 |
| N2 | 0.1 mg/ml Serotonin | pat-3(kq24, YYFF) | 0.1 mg/ml Fluoxetine | -4.04615 | 0.5053881 | -8.01 | <.0001* | -5.74818 | -2.34411 |
| N2 | 0.1 mg/ml Serotonin | pat-3(kq24, YYFF) | 0.1 mg/ml Serotonin | -4.63575 | 0.5029742 | -9.22 | <.0001* | -6.32966 | -2.94185 |
| N2 | 0.1 mg/ml Serotonin | tln-1(kq387) | 0.1 mg/ml Fluoxetine | -3.53714 | 0.5089633 | -6.95 | <.0001* | -5.25121 | -1.82307 |
| N2 | 0.1 mg/ml Serotonin | tln-1(kq387) | 0.1 mg/ml Serotonin | -4.14493 | 0.5048794 | -8.21 | <.0001* | -5.84525 | -2.44461 |
| N2 | 0.1 mg/ml Serotonin | mec-4(u253) | 0.1 mg/ml Serotonin | -3.93041 | 0.5060511 | -7.77 | <.0001* | -5.63468 | -2.22614 |
| pat-3(kq24, YYFF) | 0.1 mg/ml Fluoxetine | pat-3(kq24, YYFF) | 0.1 mg/ml Serotonin | -0.58961 | 0.0921672 | -6.4 | <.0001* | -0.9 | -0.27921 |
| pat-3(kq24, YYFF) | 0.1 mg/ml Fluoxetine | mec-4(u253) | 0.1 mg/ml Fluoxetine | 3.8236 | 0.5055365 | 7.56 | <.0001* | 2.12107 | 5.52613 |
| pat-3(kq24, YYFF) | 0.1 mg/ml Serotonin | tln-1(kq387) | 0.1 mg/ml Fluoxetine | 1.09861 | 0.1100964 | 9.98 | <.0001* | 0.72783 | 1.46939 |
| pat-3(kq24, YYFF) | 0.1 mg/ml Serotonin | tln-1(kq387) | 0.1 mg/ml Serotonin | 0.49082 | 0.0893353 | 5.49 | <.0001* | 0.18996 | 0.79169 |
| pat-3(kq24, YYFF) | 0.1 mg/ml Serotonin | mec-4(u253) | 0.1 mg/ml Fluoxetine | 4.41321 | 0.5031232 | 8.77 | <.0001* | 2.7188 | 6.10761 |
| pat-3(kq24, YYFF) | 0.1 mg/ml Serotonin | mec-4(u253) | 0.1 mg/ml Serotonin | 0.70534 | 0.0957354 | 7.37 | <.0001* | 0.38293 | 1.02776 |
| pat-3(kq7923, Y792F) | 0.1 mg/ml Fluoxetine | pat-3(kq7923, Y792F) | 0.1 mg/ml Serotonin | -0.76703 | 0.0847508 | -9.05 | <.0001* | -1.05245 | -0.4816 |
| pat-3(kq7923, Y792F) | 0.1 mg/ml Fluoxetine | pat-3(kq8042, Y804E) | 0.1 mg/ml Fluoxetine | 0.86043 | 0.141336 | 6.09 | <.0001* | 0.38444 | 1.33642 |
| pat-3(kq7923, Y792F) | 0.1 mg/ml Fluoxetine | pat-3(kq8042, Y804E) | 0.1 mg/ml Serotonin | 1.23648 | 0.1636754 | 7.55 | <.0001* | 0.68526 | 1.7877 |
| pat-3(kq7923, Y792F) | 0.1 mg/ml Fluoxetine | pat-3(kq8043, Y804F) | 0.1 mg/ml Fluoxetine | 1.19392 | 0.1608845 | 7.42 | <.0001* | 0.6521 | 1.73575 |
| pat-3(kq7923, Y792F) | 0.1 mg/ml Fluoxetine | pat-3(kq8043, Y804F) | 0.1 mg/ml Serotonin | 1.35155 | 0.1715831 | 7.88 | <.0001* | 0.7737 | 1.9294 |
| pat-3(kq7923, Y792F) | 0.1 mg/ml Fluoxetine | pat-3(kq24, YYFF) | 0.1 mg/ml Serotonin | -0.73397 | 0.0898933 | -8.16 | <.0001* | -1.03671 | -0.43123 |
| pat-3(kq7923, Y792F) | 0.1 mg/ml Fluoxetine | mec-4(u253) | 0.1 mg/ml Fluoxetine | 3.67924 | 0.5051269 | 7.28 | <.0001* | 1.97809 | 5.38039 |

| | | | | | | | | | |
|---|---|---|---|---|---|---|---|---|---|
| pat-3(kq7923, Y792F) | 0.1 mg/ml Serotonin | pat-3(kq8042, Y804E) | 0.1 mg/ml Fluoxetine | 1.62746 | 0.1306046 | 12.46 | <.0001* | 1.18761 | 2.0673 |
| pat-3(kq7923, Y792F) | 0.1 mg/ml Serotonin | pat-3(kq8042, Y804E) | 0.1 mg/ml Serotonin | 2.00351 | 0.1545035 | 12.97 | <.0001* | 1.48317 | 2.52384 |
| pat-3(kq7923, Y792F) | 0.1 mg/ml Serotonin | pat-3(kq8043, Y804F) | 0.1 mg/ml Fluoxetine | 1.96095 | 0.1515438 | 12.94 | <.0001* | 1.45058 | 2.47131 |
| pat-3(kq7923, Y792F) | 0.1 mg/ml Serotonin | pat-3(kq8043, Y804F) | 0.1 mg/ml Serotonin | 2.11858 | 0.1628571 | 13.01 | <.0001* | 1.57011 | 2.66704 |
| pat-3(kq7923, Y792F) | 0.1 mg/ml Serotonin | pat-3(kq24, YYFF) | 0.1 mg/ml Fluoxetine | 0.62266 | 0.0871589 | 7.14 | <.0001* | 0.32913 | 0.91619 |
| pat-3(kq7923, Y792F) | 0.1 mg/ml Serotonin | tln-1(kq387) | 0.1 mg/ml Fluoxetine | 1.13167 | 0.1059392 | 10.68 | <.0001* | 0.77489 | 1.48845 |
| pat-3(kq7923, Y792F) | 0.1 mg/ml Serotonin | tln-1(kq387) | 0.1 mg/ml Serotonin | 0.52388 | 0.0841587 | 6.22 | <.0001* | 0.24045 | 0.80731 |
| pat-3(kq7923, Y792F) | 0.1 mg/ml Serotonin | mec-4(u253) | 0.1 mg/ml Fluoxetine | 4.44627 | 0.5022299 | 8.85 | <.0001* | 2.75487 | 6.13766 |
| pat-3(kq7923, Y792F) | 0.1 mg/ml Serotonin | mec-4(u253) | 0.1 mg/ml Serotonin | 0.7384 | 0.090924 | 8.12 | <.0001* | 0.43219 | 1.04461 |
| pat-3(kq8042, Y804E) | 0.1 mg/ml Fluoxetine | pat-3(kq24, YYFF) | 0.1 mg/ml Fluoxetine | -1.00479 | 0.142793 | -7.04 | <.0001* | -1.48569 | -0.5239 |
| pat-3(kq8042, Y804E) | 0.1 mg/ml Fluoxetine | pat-3(kq24, YYFF) | 0.1 mg/ml Serotonin | -1.5944 | 0.1339988 | -11.9 | <.0001* | -2.04568 | -1.14312 |
| pat-3(kq8042, Y804E) | 0.1 mg/ml Fluoxetine | tln-1(kq387) | 0.1 mg/ml Serotonin | -1.10358 | 0.1409818 | -7.83 | <.0001* | -1.57837 | -0.62878 |
| pat-3(kq8042, Y804E) | 0.1 mg/ml Fluoxetine | mec-4(u253) | 0.1 mg/ml Fluoxetine | 2.81881 | 0.5148087 | 5.48 | <.0001* | 1.08505 | 4.55257 |
| pat-3(kq8042, Y804E) | 0.1 mg/ml Fluoxetine | mec-4(u253) | 0.1 mg/ml Serotonin | -0.88906 | 0.1451218 | -6.13 | <.0001* | -1.3778 | -0.40032 |
| pat-3(kq8042, Y804E) | 0.1 mg/ml Serotonin | pat-3(kq24, YYFF) | 0.1 mg/ml Fluoxetine | -1.38084 | 0.1649352 | -8.37 | <.0001* | -1.93631 | -0.82538 |
| pat-3(kq8042, Y804E) | 0.1 mg/ml Serotonin | pat-3(kq24, YYFF) | 0.1 mg/ml Serotonin | -1.97045 | 0.1573831 | -12.52 | <.0001* | -2.50048 | -1.44042 |
| pat-3(kq8042, Y804E) | 0.1 mg/ml Serotonin | tln-1(kq387) | 0.1 mg/ml Fluoxetine | -0.87184 | 0.1755848 | -4.97 | <.0001* | -1.46317 | -0.28051 |
| pat-3(kq8042, Y804E) | 0.1 mg/ml Serotonin | tln-1(kq387) | 0.1 mg/ml Serotonin | -1.47963 | 0.1633696 | -9.06 | <.0001* | -2.02982 | -0.92943 |
| pat-3(kq8042, Y804E) | 0.1 mg/ml Serotonin | mec-4(u253) | 0.1 mg/ml Serotonin | -1.26511 | 0.1669554 | -7.58 | <.0001* | -1.82738 | -0.70284 |
| pat-3(kq8043, Y804F) | 0.1 mg/ml Fluoxetine | pat-3(kq24, YYFF) | 0.1 mg/ml Fluoxetine | -1.33829 | 0.162166 | -8.25 | <.0001* | -1.88442 | -0.79215 |
| pat-3(kq8043, Y804F) | 0.1 mg/ml Fluoxetine | pat-3(kq24, YYFF) | 0.1 mg/ml Serotonin | -1.92789 | 0.1544786 | -12.48 | <.0001* | -2.44814 | -1.40764 |
| pat-3(kq8043, Y804F) | 0.1 mg/ml Fluoxetine | tln-1(kq387) | 0.1 mg/ml Serotonin | -1.43707 | 0.1605734 | -8.95 | <.0001* | -1.97784 | -0.89629 |
| pat-3(kq8043, Y804F) | 0.1 mg/ml Fluoxetine | mec-4(u253) | 0.1 mg/ml Serotonin | -1.22255 | 0.1642203 | -7.44 | <.0001* | -1.77561 | -0.66949 |

| | | | | | | | | | |
|---|---|---|---|---|---|---|---|---|---|
| pat-3(kq8043, Y804F) | 0.1 mg/ml Serotonin | pat-3(kq24, YYFF) | 0.1 mg/ml Fluoxetine | -1.49591 | 0.1727852 | -8.66 | <.0001* | -2.07782 | -0.91401 |
| pat-3(kq8043, Y804F) | 0.1 mg/ml Serotonin | pat-3(kq24, YYFF) | 0.1 mg/ml Serotonin | -2.08552 | 0.1655915 | -12.59 | <.0001* | -2.6432 | -1.52785 |
| pat-3(kq8043, Y804F) | 0.1 mg/ml Serotonin | tln-1(kq387) | 0.1 mg/ml Fluoxetine | -0.98691 | 0.1829786 | -5.39 | <.0001* | -1.60314 | -0.37068 |
| pat-3(kq8043, Y804F) | 0.1 mg/ml Serotonin | tln-1(kq387) | 0.1 mg/ml Serotonin | -1.5947 | 0.1712914 | -9.31 | <.0001* | -2.17157 | -1.01782 |
| pat-3(kq8043, Y804F) | 0.1 mg/ml Serotonin | mec-4(u253) | 0.1 mg/ml Serotonin | -1.38018 | 0.1747147 | -7.9 | <.0001* | -1.96858 | -0.79178 |
| tln-1(kq387) | 0.1 mg/ml Fluoxetine | tln-1(kq387) | 0.1 mg/ml Serotonin | -0.60779 | 0.1184964 | -5.13 | <.0001* | -1.00686 | -0.20872 |
| tln-1(kq387) | 0.1 mg/ml Fluoxetine | mec-4(u253) | 0.1 mg/ml Fluoxetine | 3.3146 | 0.5091106 | 6.51 | <.0001* | 1.60003 | 5.02917 |
| tln-1(kq387) | 0.1 mg/ml Serotonin | mec-4(u253) | 0.1 mg/ml Fluoxetine | 3.92238 | 0.5050279 | 7.77 | <.0001* | 2.22156 | 5.6232 |
| N2 | 0.1 mg/ml Fluoxetine | pat-3(kq8043, Y804F) | 0.1 mg/ml Serotonin | -1.99058 | 0.4089188 | -4.87 | 0.0001* | -3.36772 | -0.61343 |
| N2 | 0.1 mg/ml Serotonin | pat-3(kq8043, Y804F) | 0.1 mg/ml Serotonin | -2.55023 | 0.5237776 | -4.87 | 0.0001* | -4.3142 | -0.78627 |
| pat-3(kq8043, Y804F) | 0.1 mg/ml Fluoxetine | tln-1(kq387) | 0.1 mg/ml Fluoxetine | -0.82928 | 0.1729862 | -4.79 | 0.0002* | -1.41186 | -0.2467 |
| pat-3(kq8043, Y804F) | 0.1 mg/ml Fluoxetine | mec-4(u253) | 0.1 mg/ml Fluoxetine | 2.48532 | 0.5205151 | 4.77 | 0.0002* | 0.73234 | 4.23829 |
| pat-3(kq8042, Y804E) | 0.1 mg/ml Serotonin | mec-4(u253) | 0.1 mg/ml Fluoxetine | 2.44276 | 0.5213845 | 4.69 | 0.0003* | 0.68685 | 4.19866 |
| pat-3(kq8043, Y804F) | 0.1 mg/ml Serotonin | mec-4(u253) | 0.1 mg/ml Fluoxetine | 2.32769 | 0.5239207 | 4.44 | 0.0009* | 0.56324 | 4.09213 |
| pat-3(kq24, YYFF) | 0.1 mg/ml Fluoxetine | tln-1(kq387) | 0.1 mg/ml Fluoxetine | 0.50901 | 0.1206457 | 4.22 | 0.0023* | 0.1027 | 0.91531 |
