## Supplementary Table 5 for "The β integrin modulates serotonin sensitivity via NPxY motifs to regulate egg laying and mechanosensation behaviors in *C. elegans*"

| Level | - Level | Difference | Std Err Dif | Lower CL | Upper CL | p-Value |
|---|---|---|---|---|---|---|
| mec-4(u253) | pat-3(kq7923, Y792F) | 0.085714 | 0.6690899 | -1.8993 | 2.070732 | 1 |
| N2 | pat-3(kq8041, Y804A) | 1.6 | 0.5729744 | -0.09987 | 3.299868 | 0.0802 |
| N2 | pat-3(kq8043, Y804F) | 0.228571 | 0.5729744 | -1.4713 | 1.928439 | 0.9997 |
| N2 | pat-3(kq8042, Y804E) | 0.129412 | 0.4434283 | -1.18613 | 1.44495 | 0.9999 |
| pat-3(kq8041, Y804A) | pat-3(kq7923, Y792F) | 0.657143 | 0.6690899 | -1.32787 | 2.642161 | 0.9576 |
| pat-3(kq8041, Y804A) | mec-4(u253) | 0.571429 | 0.6690899 | -1.41359 | 2.556446 | 0.9788 |
| pat-3(kq8042, Y804E) | pat-3(kq8041, Y804A) | 1.470588 | 0.562148 | -0.19716 | 3.138337 | 0.1245 |
| pat-3(kq8042, Y804E) | pat-3(kq8043, Y804F) | 0.09916 | 0.562148 | -1.56859 | 1.766908 | 1 |
| pat-3(kq8043, Y804F) | mec-4(u253) | 1.942857 | 0.6690899 | -0.04216 | 3.927875 | 0.0596 |
| pat-3(kq8043, Y804F) | pat-3(kq8041, Y804A) | 1.371429 | 0.6690899 | -0.61359 | 3.356446 | 0.3855 |
