## Supplementary Table 6 for "The β integrin modulates serotonin sensitivity via NPxY motifs to regulate egg laying and mechanosensation behaviors in *C. elegans*"

| Strains | Sections | /Strains | /Sections | Odds Ratio | Std Error | t Ratio | Prob>\|t\| | Lower 95% | Upper 95% |
|---|---|---|---|---|---|---|---|---|---|
| N2 | Anterior | pat-3(kq7923, Y792F) RS | Posterior | 6.2327 | 3.4252 | 3.33 | 0.1131 | 0.86442 | 44.94 |
| N2 | Anterior | pat-3(kq8041, Y804A) | Posterior | 5.3491 | 2.9709 | 3.02 | 0.2519 | 0.72643 | 39.388 |
| N2 | Anterior | pat-3(kq8041, Y804A) | Anterior | 5.0816 | 2.8343 | 2.91 | 0.3169 | 0.68427 | 37.738 |
| N2 | Anterior | pat-3(kq8043, Y804F) | Posterior | 4.7727 | 2.6743 | 2.79 | 0.4053 | 0.63673 | 35.774 |
| N2 | Anterior | N2, 5-HT | Anterior | 4.1882 | 2.3725 | 2.53 | 0.61 | 0.54661 | 32.091 |
| N2 | Anterior | pat-3(kq7923, Y792F) RS | Anterior | 3.9264 | 2.2386 | 2.4 | 0.7094 | 0.50574 | 30.484 |
| N2 | Anterior | pat-3(kq24, YYFF) RS | Anterior | 3.1016 | 1.8129 | 1.94 | 0.9472 | 0.37935 | 25.358 |
| N2 | Anterior | N2 | Posterior | 0.7472 | 0.5746 | -0.38 | 1 | 0.04707 | 11.861 |
| N2 | Anterior | pat-3(kq7923, Y792F) | Anterior | 1.5126 | 0.986 | 0.63 | 1 | 0.14524 | 15.753 |
| N2 | Anterior | pat-3(kq8042, Y804E) | Anterior | 2.0335 | 1.2589 | 1.15 | 1 | 0.21966 | 18.825 |
| N2 | Anterior | pat-3(kq8043, Y804F) | Anterior | 0.7472 | 0.5746 | -0.38 | 1 | 0.04707 | 11.861 |
| N2 | Posterior | pat-3(kq7923, Y792F) RS | Posterior | 8.3417 | 5.1775 | 3.42 | 0.0874 | 0.89588 | 77.671 |
| N2 | Posterior | pat-3(kq8041, Y804A) | Posterior | 7.159 | 4.4806 | 3.15 | 0.186 | 0.75466 | 67.913 |
| N2 | Posterior | pat-3(kq8041, Y804A) | Anterior | 6.8011 | 4.2708 | 3.05 | 0.233 | 0.71154 | 65.006 |
| N2 | Posterior | pat-3(kq8043, Y804F) | Posterior | 6.3876 | 4.0258 | 2.94 | 0.2989 | 0.66279 | 61.559 |
| N2 | Posterior | N2, 5-HT | Anterior | 5.6054 | 3.5633 | 2.71 | 0.4646 | 0.57036 | 55.088 |
| N2 | Posterior | pat-3(kq7923, Y792F) RS | Anterior | 5.255 | 3.3578 | 2.6 | 0.5556 | 0.52847 | 52.255 |
| N2 | Posterior | pat-3(kq24, YYFF) RS | Anterior | 4.151 | 2.7058 | 2.18 | 0.8482 | 0.39857 | 43.232 |
| N2 | Posterior | pat-3(kq8042, Y804E) | Anterior | 2.7216 | 1.8589 | 1.47 | 0.9981 | 0.23363 | 31.705 |
| N2 | Posterior | pat-3(kq7923, Y792F) | Anterior | 2.0244 | 1.4431 | 0.99 | 1 | 0.15611 | 26.253 |
| N2 | Posterior | pat-3(kq8043, Y804F) | Anterior | 1 | 0.8214 | 0 | 1 | 0.05219 | 19.16 |
| N2, 5-HT | Anterior | pat-3(kq24, YYFF) RS | Posterior | 2.2336 | 0.7122 | 2.52 | 0.6164 | 0.70993 | 7.027 |
| N2, 5-HT | Anterior | pat-3(kq7923, Y792F) RS | Posterior | 1.4882 | 0.5039 | 1.17 | 0.9999 | 0.44061 | 5.026 |

| | | | | | | | | | |
|---|---|---|---|---|---|---|---|---|---|
| N2, 5-HT | Anterior | pat-3(kq24, YYFF) RS | Anterior | 0.7405 | 0.2909 | -0.76 | 1 | 0.18043 | 3.039 |
| N2, 5-HT | Anterior | pat-3(kq7923, Y792F) RS | Anterior | 0.9375 | 0.3479 | -0.17 | 1 | 0.247 | 3.558 |
| N2, 5-HT | Posterior | pat-3(kq7923, Y792F) RS | Posterior | 0.4033 | 0.1089 | -3.36 | 0.1026 | 0.15277 | 1.064 |
| N2, 5-HT | Posterior | pat-3(kq24, YYFF) RS | Posterior | 0.6053 | 0.1482 | -2.05 | 0.9098 | 0.25103 | 1.459 |
| pat-3(kq24, YYFF) | Anterior | N2, 5-HT | Posterior | 2.1647 | 0.5643 | 2.96 | 0.286 | 0.84807 | 5.525 |
| pat-3(kq24, YYFF) | Anterior | pat-3(kq24, YYFF) RS | Anterior | 0.4344 | 0.1569 | -2.31 | 0.7728 | 0.11857 | 1.591 |
| pat-3(kq24, YYFF) | Anterior | pat-3(kq7923, Y792F) RS | Anterior | 0.5499 | 0.1855 | -1.77 | 0.9792 | 0.16351 | 1.85 |
| pat-3(kq24, YYFF) | Anterior | N2, 5-HT | Anterior | 0.5866 | 0.1943 | -1.61 | 0.9934 | 0.17836 | 1.929 |
| pat-3(kq24, YYFF) | Anterior | pat-3(kq24, YYFF) RS | Posterior | 1.3102 | 0.3655 | 0.97 | 1 | 0.48058 | 3.572 |
| pat-3(kq24, YYFF) | Anterior | pat-3(kq7923, Y792F) RS | Posterior | 0.8729 | 0.2631 | -0.45 | 1 | 0.29547 | 2.579 |
| pat-3(kq24, YYFF) | Posterior | mec-4(u253) | Posterior | 0.6729 | 0.1613 | -1.65 | 0.9909 | 0.28428 | 1.593 |
| pat-3(kq24, YYFF) RS | Anterior | pat-3(kq24, YYFF) RS | Posterior | 3.0161 | 1.0554 | 3.15 | 0.1814 | 0.85734 | 10.611 |
| pat-3(kq24, YYFF) RS | Anterior | pat-3(kq7923, Y792F) RS | Posterior | 2.0095 | 0.7395 | 1.9 | 0.9571 | 0.53529 | 7.544 |
| pat-3(kq24, YYFF) RS | Anterior | pat-3(kq7923, Y792F) RS | Anterior | 1.266 | 0.5039 | 0.59 | 1 | 0.30266 | 5.295 |
| pat-3(kq24, YYFF) RS | Posterior | pat-3(kq7923, Y792F) RS | Anterior | 0.4197 | 0.1365 | -2.67 | 0.4983 | 0.13034 | 1.352 |
| pat-3(kq24, YYFF) RS | Posterior | pat-3(kq7923, Y792F) RS | Posterior | 0.6663 | 0.1917 | -1.41 | 0.9989 | 0.23683 | 1.874 |
| pat-3(kq7921, Y792A) | Anterior | pat-3(kq7921, Y792A) | Posterior | 0.5601 | 0.1043 | -3.11 | 0.2012 | 0.28684 | 1.094 |
| pat-3(kq7921, Y792A) | Anterior | mec-4(u253) | Posterior | 1.848 | 0.3854 | 2.94 | 0.2977 | 0.87308 | 3.911 |
| pat-3(kq7921, Y792A) | Anterior | mec-4(u253) | Anterior | 0.5976 | 0.1114 | -2.76 | 0.4267 | 0.3057 | 1.168 |
| pat-3(kq7921, Y792A) | Anterior | pat-3(kq7923, Y792F) | Posterior | 0.913 | 0.1743 | -0.48 | 1 | 0.45968 | 1.813 |
| pat-3(kq7921, Y792A) | Anterior | pat-3(kq8042, Y804E) | Posterior | 0.913 | 0.1743 | -0.48 | 1 | 0.45968 | 1.813 |
| pat-3(kq7921, Y792A) | Posterior | pat-3(kq7923, Y792F) | Posterior | 1.63 | 0.3009 | 2.65 | 0.5157 | 0.83944 | 3.165 |

| | | | | | | | | | |
|---|---|---|---|---|---|---|---|---|---|
| pat-3(kq7921, Y792A) | Posterior | pat-3(kq8042, Y804E) | Posterior | 1.63 | 0.3009 | 2.65 | 0.5157 | 0.83944 | 3.165 |
| pat-3(kq7921, Y792A) | Posterior | mec-4(u253) | Anterior | 1.067 | 0.1921 | 0.36 | 1 | 0.55856 | 2.038 |
| pat-3(kq7923, Y792F) | Anterior | pat-3(kq24, YYFF) | Anterior | 4.7204 | 2.1821 | 3.36 | 0.1045 | 0.89594 | 24.87 |
| pat-3(kq7923, Y792F) | Anterior | pat-3(kq7923, Y792F) RS | Posterior | 4.1205 | 1.9268 | 3.03 | 0.2468 | 0.76724 | 22.129 |
| pat-3(kq7923, Y792F) | Anterior | pat-3(kq8041, Y804A) | Posterior | 3.5363 | 1.6779 | 2.66 | 0.5034 | 0.64242 | 19.466 |
| pat-3(kq7923, Y792F) | Anterior | pat-3(kq8041, Y804A) | Anterior | 3.3595 | 1.6033 | 2.54 | 0.6014 | 0.60427 | 18.678 |
| pat-3(kq7923, Y792F) | Anterior | pat-3(kq8043, Y804F) | Posterior | 3.1553 | 1.5153 | 2.39 | 0.714 | 0.56141 | 17.733 |
| pat-3(kq7923, Y792F) | Anterior | N2, 5-HT | Anterior | 2.7689 | 1.3494 | 2.09 | 0.8938 | 0.48021 | 15.965 |
| pat-3(kq7923, Y792F) | Anterior | pat-3(kq7923, Y792F) RS | Anterior | 2.5958 | 1.2762 | 1.94 | 0.9462 | 0.44336 | 15.198 |
| pat-3(kq7923, Y792F) | Anterior | pat-3(kq24, YYFF) RS | Anterior | 2.0505 | 1.0421 | 1.41 | 0.9989 | 0.32991 | 12.744 |
| pat-3(kq7923, Y792F) | Anterior | pat-3(kq8042, Y804E) | Anterior | 1.3444 | 0.7362 | 0.54 | 1 | 0.18774 | 9.627 |
| pat-3(kq7923, Y792F) | Anterior | pat-3(kq8043, Y804F) | Anterior | 0.494 | 0.3521 | -0.99 | 1 | 0.03809 | 6.406 |
| pat-3(kq7923, Y792F) | Posterior | mec-4(u253) | Posterior | 2.0241 | 0.4194 | 3.4 | 0.0912 | 0.96109 | 4.263 |
| pat-3(kq7923, Y792F) | Posterior | mec-4(u253) | Anterior | 0.6546 | 0.121 | -2.29 | 0.7839 | 0.33672 | 1.273 |
| pat-3(kq7923, Y792F) | Posterior | pat-3(kq8042, Y804E) | Posterior | 1 | 0.1894 | 0 | 1 | 0.50626 | 1.975 |
| pat-3(kq7923, Y792F) RS | Anterior | pat-3(kq7923, Y792F) RS | Posterior | 1.5874 | 0.5471 | 1.34 | 0.9995 | 0.4598 | 5.48 |
| pat-3(kq8041, Y804A) | Anterior | pat-3(kq8043, Y804F) | Anterior | 0.147 | 0.0923 | -3.05 | 0.233 | 0.01538 | 1.405 |
| pat-3(kq8041, Y804A) | Anterior | pat-3(kq8042, Y804E) | Anterior | 0.4002 | 0.1726 | -2.12 | 0.8788 | 0.08487 | 1.887 |
| pat-3(kq8041, Y804A) | Anterior | pat-3(kq24, YYFF) RS | Posterior | 1.8409 | 0.558 | 2.01 | 0.9237 | 0.61912 | 5.474 |
| pat-3(kq8041, Y804A) | Anterior | pat-3(kq24, YYFF) RS | Anterior | 0.6103 | 0.232 | -1.3 | 0.9997 | 0.15563 | 2.394 |
| pat-3(kq8041, Y804A) | Anterior | pat-3(kq8041, Y804A) | Posterior | 1.0526 | 0.3512 | 0.15 | 1 | 0.31723 | 3.493 |
| pat-3(kq8041, Y804A) | Anterior | pat-3(kq8043, Y804F) | Posterior | 0.9392 | 0.321 | -0.18 | 1 | 0.27485 | 3.209 |

| | | | | | | | | | |
|---|---|---|---|---|---|---|---|---|---|
| pat-3(kq8041, Y804A) | Anterior | pat-3(kq24, YYFF) | Anterior | 1.4051 | 0.4441 | 1.08 | 1 | 0.45107 | 4.377 |
| pat-3(kq8041, Y804A) | Anterior | N2, 5-HT | Anterior | 0.8242 | 0.2899 | -0.55 | 1 | 0.23274 | 2.919 |
| pat-3(kq8041, Y804A) | Anterior | pat-3(kq7923, Y792F) RS | Anterior | 0.7727 | 0.2763 | -0.72 | 1 | 0.21363 | 2.795 |
| pat-3(kq8041, Y804A) | Anterior | pat-3(kq7923, Y792F) RS | Posterior | 1.2265 | 0.3972 | 0.63 | 1 | 0.38293 | 3.929 |
| pat-3(kq8041, Y804A) | Posterior | pat-3(kq8043, Y804F) | Anterior | 0.1397 | 0.0874 | -3.15 | 0.186 | 0.01472 | 1.325 |
| pat-3(kq8041, Y804A) | Posterior | pat-3(kq8042, Y804E) | Anterior | 0.3802 | 0.1628 | -2.26 | 0.8053 | 0.08152 | 1.773 |
| pat-3(kq8041, Y804A) | Posterior | pat-3(kq24, YYFF) RS | Posterior | 1.7488 | 0.5225 | 1.87 | 0.9627 | 0.59748 | 5.119 |
| pat-3(kq8041, Y804A) | Posterior | pat-3(kq24, YYFF) RS | Anterior | 0.5798 | 0.2184 | -1.45 | 0.9984 | 0.14971 | 2.246 |
| pat-3(kq8041, Y804A) | Posterior | pat-3(kq8043, Y804F) | Posterior | 0.8922 | 0.3015 | -0.34 | 1 | 0.26477 | 3.007 |
| pat-3(kq8041, Y804A) | Posterior | pat-3(kq24, YYFF) | Anterior | 1.3348 | 0.4163 | 0.93 | 1 | 0.43502 | 4.096 |
| pat-3(kq8041, Y804A) | Posterior | N2, 5-HT | Anterior | 0.783 | 0.2725 | -0.7 | 1 | 0.22411 | 2.736 |
| pat-3(kq8041, Y804A) | Posterior | pat-3(kq7923, Y792F) RS | Anterior | 0.734 | 0.2598 | -0.87 | 1 | 0.20567 | 2.62 |
| pat-3(kq8041, Y804A) | Posterior | pat-3(kq7923, Y792F) RS | Posterior | 1.1652 | 0.3726 | 0.48 | 1 | 0.36917 | 3.678 |
| pat-3(kq8042, Y804E) | Anterior | pat-3(kq24, YYFF) | Anterior | 3.5112 | 1.4564 | 3.03 | 0.247 | 0.79044 | 15.597 |
| pat-3(kq8042, Y804E) | Anterior | pat-3(kq7923, Y792F) RS | Posterior | 3.065 | 1.2895 | 2.66 | 0.5034 | 0.67544 | 13.908 |
| pat-3(kq8042, Y804E) | Anterior | pat-3(kq8043, Y804F) | Posterior | 2.347 | 1.0203 | 1.96 | 0.9399 | 0.49181 | 11.2 |
| pat-3(kq8042, Y804E) | Anterior | N2, 5-HT | Anterior | 2.0596 | 0.9116 | 1.63 | 0.9922 | 0.41957 | 10.11 |
| pat-3(kq8042, Y804E) | Anterior | pat-3(kq7923, Y792F) RS | Anterior | 1.9309 | 0.8636 | 1.47 | 0.998 | 0.38678 | 9.639 |
| pat-3(kq8042, Y804E) | Anterior | pat-3(kq8043, Y804F) | Anterior | 0.3674 | 0.251 | -1.47 | 0.9981 | 0.03154 | 4.28 |
| pat-3(kq8042, Y804E) | Anterior | pat-3(kq24, YYFF) RS | Anterior | 1.5252 | 0.7099 | 0.91 | 1 | 0.28618 | 8.129 |
| pat-3(kq8042, Y804E) | Posterior | mec-4(u253) | Posterior | 2.0241 | 0.4194 | 3.4 | 0.0912 | 0.96109 | 4.263 |
| pat-3(kq8042, Y804E) | Posterior | mec-4(u253) | Anterior | 0.6546 | 0.121 | -2.29 | 0.7839 | 0.33672 | 1.273 |

| | | | | | | | | | |
|---|---|---|---|---|---|---|---|---|---|
| pat-3(kq8043, Y804F) | Anterior | pat-3(kq7923, Y792F) RS | Posterior | 8.3417 | 5.1775 | 3.42 | 0.0874 | 0.89588 | 77.671 |
| pat-3(kq8043, Y804F) | Anterior | pat-3(kq8043, Y804F) | Posterior | 6.3876 | 4.0258 | 2.94 | 0.2989 | 0.66279 | 61.559 |
| pat-3(kq8043, Y804F) | Anterior | N2, 5-HT | Anterior | 5.6054 | 3.5633 | 2.71 | 0.4646 | 0.57036 | 55.088 |
| pat-3(kq8043, Y804F) | Anterior | pat-3(kq7923, Y792F) RS | Anterior | 5.255 | 3.3578 | 2.6 | 0.5556 | 0.52847 | 52.255 |
| pat-3(kq8043, Y804F) | Anterior | pat-3(kq24, YYFF) RS | Anterior | 4.151 | 2.7058 | 2.18 | 0.8482 | 0.39857 | 43.232 |
| pat-3(kq8043, Y804F) | Posterior | pat-3(kq24, YYFF) RS | Posterior | 1.9601 | 0.6034 | 2.19 | 0.8469 | 0.6481 | 5.928 |
| pat-3(kq8043, Y804F) | Posterior | pat-3(kq24, YYFF) | Anterior | 1.496 | 0.4796 | 1.26 | 0.9998 | 0.4725 | 4.737 |
| pat-3(kq8043, Y804F) | Posterior | N2, 5-HT | Anterior | 0.8775 | 0.3123 | -0.37 | 1 | 0.24419 | 3.154 |
| pat-3(kq8043, Y804F) | Posterior | pat-3(kq24, YYFF) RS | Anterior | 0.6499 | 0.2495 | -1.12 | 1 | 0.16347 | 2.583 |
| pat-3(kq8043, Y804F) | Posterior | pat-3(kq7923, Y792F) RS | Anterior | 0.8227 | 0.2975 | -0.54 | 1 | 0.2242 | 3.019 |
| pat-3(kq8043, Y804F) | Posterior | pat-3(kq7923, Y792F) RS | Posterior | 1.3059 | 0.4287 | 0.81 | 1 | 0.40129 | 4.25 |
